# Context-dependent reshaping of defensive responses to predators in head-fixed and freely moving mice

**DOI:** 10.1101/2025.06.09.658679

**Authors:** Marti Ritter, Lara Mariel Chirich Barreira, Lara Sach, Aileen Hakus, Sanra Kumsal Öktem, Ronny Bergmann, Anne Voigt, Dietmar Schmitz, Panayiota Poirazi, Matthew E. Larkum, Robert N. S. Sachdev

## Abstract

Behavioral responses to threat — such as fleeing, freezing, or fighting—can be innate, learned, and strongly shaped by context or competing goals. Here, we asked whether exposure to an ecologically relevant predator obligatorily elicits canonical defensive behaviors across behavioral contexts. We examined predator responses in mice across four experimental conditions: one novel head-fixed reward-driven foraging task and three established paradigms in freely moving animals. In the head-fixed condition, water-deprived mice were trained to walk on a treadmill controlling a virtual environment and water reward delivery and were subsequently exposed to a live rat positioned above the lick spout. Despite the presence of the predator, most mice (5 of 7) maintained foraging performance at baseline levels. However, individual mice exhibited significant, coordinated changes in running speed, pupil diameter, eye movements, and posture, indicating engagement with the threat. To assess how context influences predator responses, we exposed 36 naïve, freely moving mice to fear-inducing stimuli, including looming visual cues, rat odor, and a live rat. Even under these conditions, defensive behaviors were variable: only a subset of mice displayed avoidance or escape, and when presented with a freely moving rat, approximately half of the mice avoided the predator. Together, these findings show that predator threat does not elicit a uniform or obligatory defensive repertoire in mice. Instead, defensive responses are expressed flexibly, and are shaped by environmental constraints, task demands, and individual variability. These results challenge the assumption that innate fear behaviors are automatically triggered by predator encounters.

**Graphical Abstract:** Using a head-fixed, reward-based foraging task and complementary freely moving paradigms, we show that mouse responses to predator-related stimuli are variable and context dependent, with limited expression of canonical defensive behaviors such as freezing or flight.

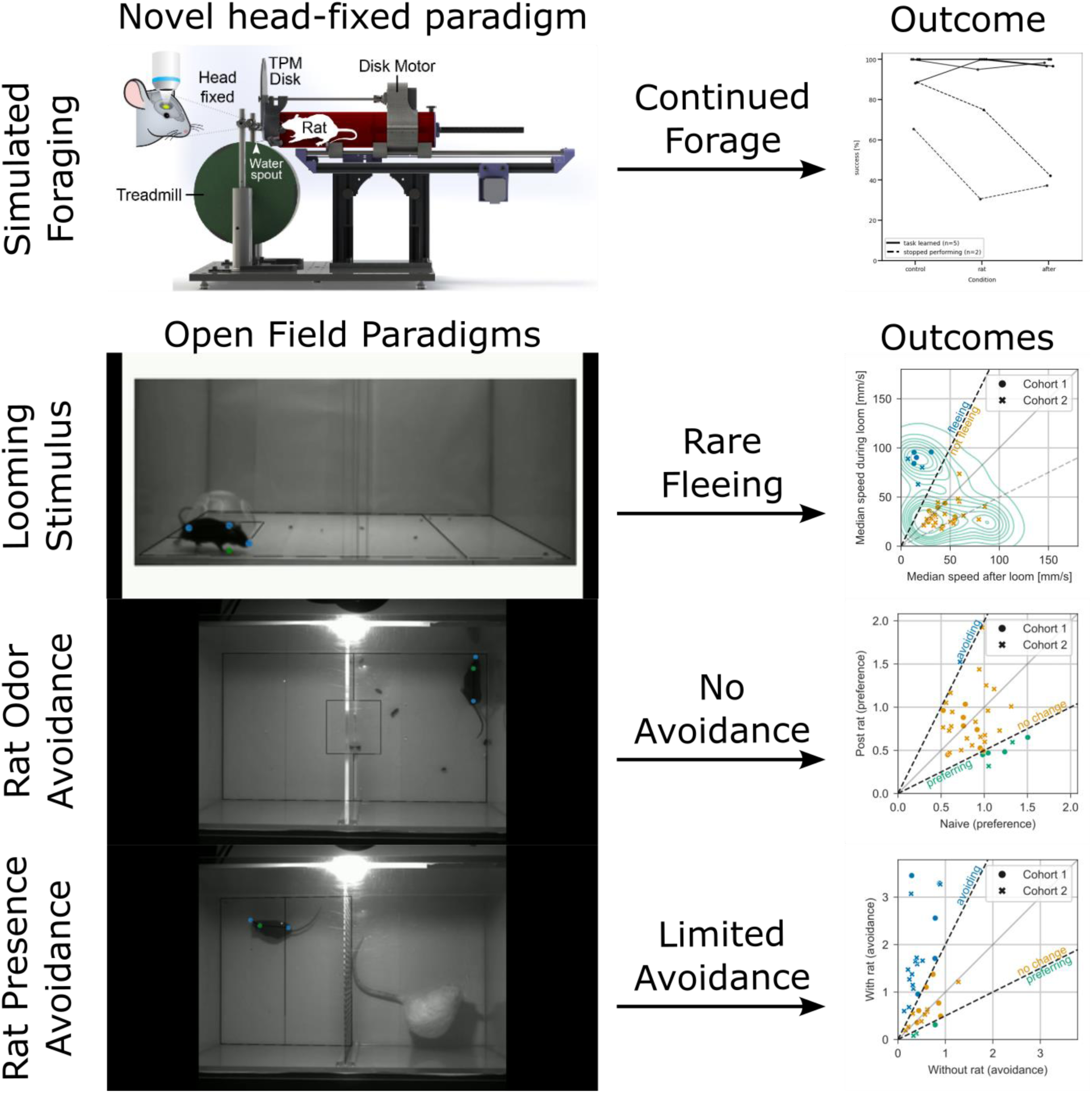

## Introduction

Threat perception and appropriate reaction to a threat are necessary for survival. Overestimating dangers, for example misinterpreting a benign sound as a predator, or underestimating a risk, for example misjudging the strength of a barrier separating prey from a predator, can lead to unwanted outcomes ranging from unnecessary anxiety to injury or death. Accordingly, threat processing has been studied extensively across species, with work traditionally focusing on three interrelated components: the detection of a threat, the behavioral and physiological responses it evokes, and the subjective experience of fear (Beckers et al. 2013; Fanselow and Pennington 2017; LeDoux and Brown 2017; Mobbs et al. 2019; Silva et al. 2016; Tovote et al. 2015). Much of what is known about threat processing derives from Pavlovian fear conditioning paradigms (McDannald 2023), in which a neutral stimulus—often an auditory cue—is paired with an aversive outcome such as a mild foot shock (Mowrer and Lamoreaux 1946). Through repeated pairing, the previously neutral cue comes to elicit defensive behaviors, including freezing, fleeing, postural changes, and accompanying physiological responses such as pupil dilation and increased heart rate (Blanchard and Blanchard 1989). These classes of paradigms offer precise experimental control and have been instrumental in delineating the neural circuits underlying threat detection and defensive behavior (Phillips and LeDoux 1992; reviewed in Tovote et al. 2015). In parallel, a range of ethologically inspired paradigms have been developed to probe so-called “innate” fear responses using evolutionarily relevant stimuli. Looming visual stimuli that mimic an approaching aerial predator can evoke rapid escape or freezing responses in rodents (Schiff 1965; Yilmaz and Meister 2013). Similarly, exposure to predator-associated odors (e.g., fox or cat urine), heights, loud sounds, or larger conspecifics has been used to elicit defensive behaviors by introducing naturalistic threats (Gibson and Walk 1960; Mongeau et al. 2003; Farmer-Dougan et al. 2005; Silva et al. 2016). These approaches are often assumed to engage hardwired defensive circuits that produce stereotyped behavioral outputs.

However, accumulating evidence suggests that defensive behaviors are not purely reflexive but are modulated by context, internal state, and available action (Maren et al. 2013). When threats occur in environments that impose physical constraints or introduce competing motivations, such as the need to obtain food or water, defensive responses may be reshaped rather than being simply expressed or suppressed (Burnett et al. 2016). Under such conditions, animals may engage in graded threat assessment or alternative coping strategies instead of canonical freezing or flight. In this study, we sought to test whether exposure to an ecologically relevant predator obligatorily elicits stereotyped defensive behaviors, or whether such responses are flexibly expressed depending on behavioral context and physical constraints. To address this, we developed a head-fixed, reward-based foraging paradigm in which mice ran on a treadmill to obtain water while exposed to a live rat positioned directly above the lick spout. The head fixed preparation is the state of the art and standard approach for circuit mapping, calcium imaging, and high-density neural recordings to understand the neural basis of sensory motor behaviors and is often used for studying fear conditioning (Lovett-Barron et al. 2014; Abs et al. 2018; Ahmad et al. 2020; Dalmay et al. 2019). Here our goal was to develop a novel paradigm that could be easily combined with state-of-the-art imaging and recording methods. To achieve this goal, we examined predator responses under conditions of constrained movement and competing motivational demands. We compared behavior in this novel context to responses observed in freely moving mice exposed to established fear-inducing stimuli, including looming visual cues, predator odor, and a live rat.

Previous work has suggested that rats can act as natural predators of mice, eliciting innate defensive responses (Karli 1956; Panksepp 1971; Blanchard et al. 1998). Here, we asked whether mice would alter their behavior in the presence of a rat when foraging for reward was required, or whether they would continue goal-directed behavior while expressing alternative, non-canonical indicators of threat engagement. Surprisingly, although head-fixed mice showed clear changes in physiological and postural measures during predator exposure, most continued to forage. Moreover, even in freely moving paradigms designed to elicit robust fear responses, mice did not consistently display avoidance, freezing, or escape. Together, these observations suggest a reassessment of how “innate” defensive behaviors are expressed across different contexts.

## Methods and Materials

All experiments were conducted in accordance with the guidelines of animal welfare of the Charité Universitätsmedizin Berlin and the local authorities, the ‘Landesamt für Gesundheit und Soziales’. Adult mice (n=43) on a C57BL / J6 background and 5 rats were used in this study. For experiments involving head fixation, seven mice (4 male, 3 female, 8-12 months old) were sourced from the internal breeding facilities. We selected naive, untreated mice available in our facility. This resulted in the given mix of ages and sexes. For experiments with freely moving mice (n=36), male mice (∼8 weeks old at start of experiments) were sourced from our in-house breeding colony, Janvier, and Charles River (n=12 each). Their origin was blinded prior to experiments, by having a separate researcher uninvolved with the experiments and analysis anonymize the mouse IDs and groups during the separation of the mice. Mice were housed socially with at least one other littermate. Five female Wistar rats served as the predator stimulus and were kept in the same facility as the mice, but in another room. For experiments with head-fixed mice, we used a different rat each day. In the freely-moving paradigms, a random rat was selected by the experimenter for each session. Both the mice and rats were kept in a 12h reversed light cycle (mice 6pm to 6am, rats 5pm to 5am). Except for mice that were actively used in the head-fixed simulated foraging paradigms and were on water restriction, all animals had ad lib access to food and water.

### Surgical procedures for head-fixation

On the day of the surgery, adult C57Bl6 mice (n=7) weighing 20-40 grams were injected with Carprofen (5 mg/kg) intraperitoneally pre-surgery, then deeply anesthetized with a mixture of Ketamine and Xylazine (Ketamine 12.5mg/ml, Xylazine 1 mg/ml, 10 µl/g dose) and placed on a heating pad maintained at 37°C. A local analgesic, Lidocaine (100 µl), was injected under the skin at the site of the incision. Lightweight aluminum headposts (3 cm long, 2mm thick, weighing 0.4 gm) were affixed to the skull using Rely X and Jet Acrylic (Ortho-Jet) black cement (Dominiak et al., 2019; Ebner et al., 2019). Post-surgery analgesia was provided over 3 days, with Carprofen injections (5 mg/kg) intraperitoneally and, if there were signs of post-operative pain, Buprenorphine (0.05-0.1 mg/kg) subcutaneously in addition.

### Simulated foraging experiment

The simulated foraging environment consisted of a rigid-foam-based circular treadmill and a 30 cm long plexiglass tube with a 7 cm diameter (**Figure 1, Video 1**). The treadmill was lightweight enough that mice could move it effortlessly. It had a rubberized surface to increase the grip of the mouse on the wheel. The movement of the treadmill was read by an encoder. The output of the encoder was used to control the position of the plexiglass tube, which was mounted on a rail and controlled by a small stepper motor (**Figure 1**). To ensure safe and stable movement, and to minimize the stress on a rat held in the tube, the top speed of the tube was limited to 4 cm per second. The output of the encoder was also linked to the PC that controlled two large monitors positioned on each side of the treadmill that were used to stream visual stimuli tethered to the movement of the treadmill.

**Figure 1.**
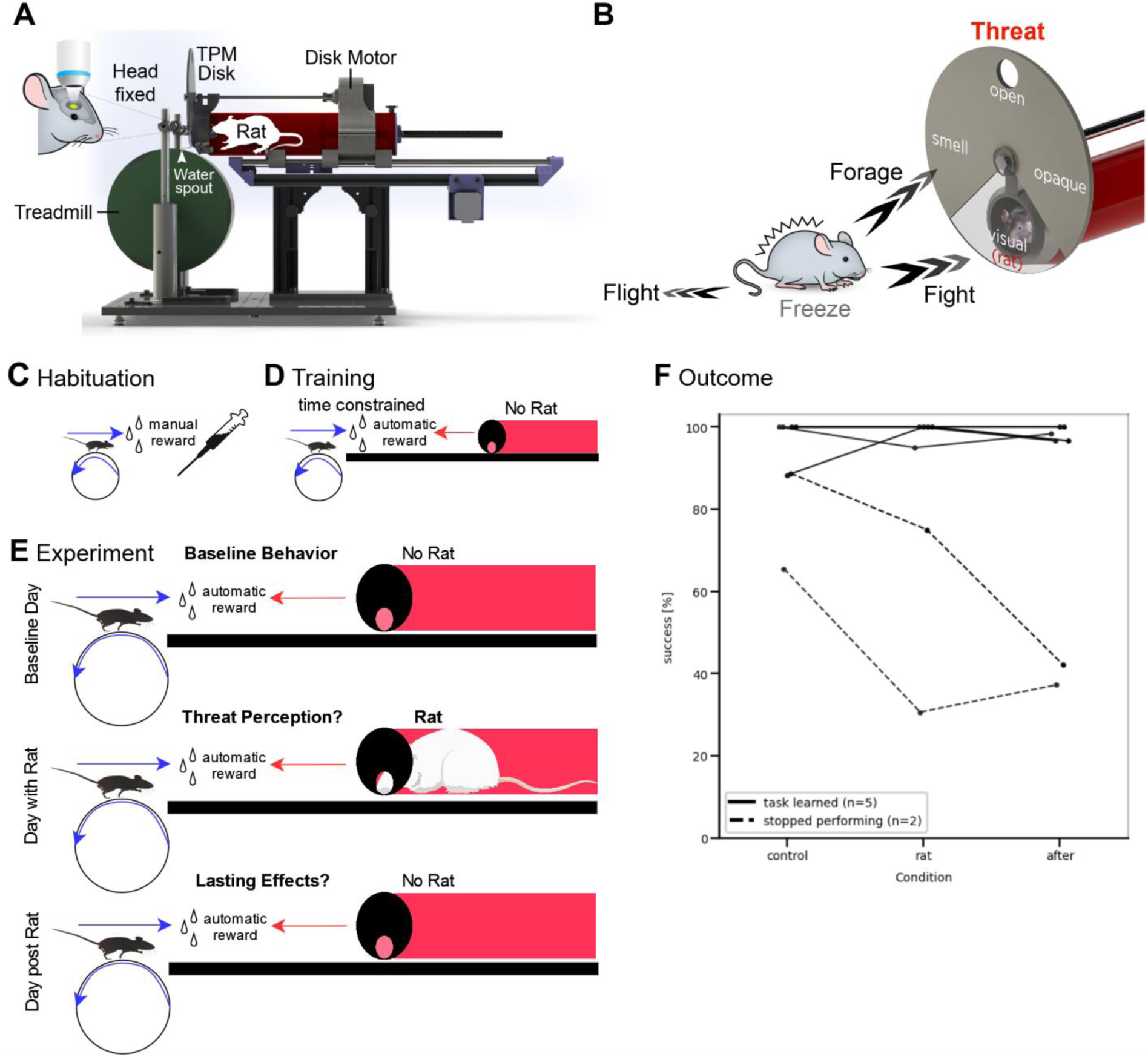
Experimental design, setup, and performance in the presence of a rat. **A)** Side view of the mechanical components in the apparatus. The movement of a treadmill was tethered to the movement of a lick spout and a large red tube. When mice walked forward, the lick tube moved towards them, when mice moved backwards, the lick tube moved away. A flashing screen (part of a virtual reality display) and a sound cue signaled the start of a trial. Mice had 12 seconds to complete a trial, if they succeeded, reward was delivered within reach of the mouse, and a new trial was initiated by resetting the position of the spout to its starting position away from the mouse. **B)** Threat perception disc. The apparatus was designed to mimic four user defined interactive conditions with the large red tube positioned over the lick spout. The full view condition was one where the opening of the tube was completely open, and it was possible for a head-fixed mouse and the rat to almost touch each other. The olfactory condition was one where odors could be delivered but visual, tactile and other elements inside the tube were blocked. The visual condition was one where mice could see inside the tube. The opaque condition blocked all cues from inside the tube. Note that auditory cues were not filtered and that a fan at the end of the tube could be activated to extract smells from the red tube. **C**) Training procedures. Habituation consisted of head fixing mice on the treadmill and manually rewarding them when they moved forward spontaneously. Following this brief, 1- to 2-day habituation period, the reward delivery was automated. Reward was dispensed after mice had moved 28.5 centimeters. This movement was sufficient to move the lick spout into a position where mice could lick the reward as it was delivered. Over days the flashing light cue of the virtual reality screens, and the sound cue were added as go-cues to begin the trial. The duration of the trial was shortened, giving mice 12 seconds to reach the spout, or move at least 28.5 cm. **D**) Data acquisition. Once all mice of the cohort reached criterion (65% success rate) baseline data was acquired. In the next session, a live rat was placed inside the tube, and the experiment was repeated, with the full interaction between the mouse and the rat. Finally, on a third day, reference data in the absence of the rat were acquired. **E**) Performance data. Of the seven mice used for these data sets, five showed no change in their performance in any of the three sessions. For the other two, performance was affected by the presence of the live rat. For both these mice, the reduction in performance persisted through the next day, post rat. Vector drawings of rat and mouse adapted from SciDraw (Branco and Costa 2020; Scidraw 2020).

The tube holding the rat was coated with a red film limiting the ability of mice to see the position of the rat, or to see whether there was a rat in the tube (**Figure 1A**). Additionally, the front of the tube was covered with a disk which could be rotated by an electronic input (**Figure 1B)**. When a rat was in the tube, the disk could be used to selectively hide or show the rat during the simulated predator encounter. The disk could be rotated to allow odors through, or to block all sensory stimuli from inside the tube. When the disk was in the open position, the rat could stick its nose out of the tube and almost touch the mouse when it was at the lick spout.

### Electronics and control of behavior

For an overview of the connections in the setup, see **Supplementary Figure 1**.

### Trial control

A finite state machine (Bpod r2, Sanworks LLC) monitored trial states (see **Supplementary Figure 2**), and the sequence of reward delivery and data acquisition. This state machine also controlled the high-resolution acquisition of movement traces from the encoder in the treadmill and triggered the cameras (recording at 100Hz).

### Motor control

In addition to the state machine, a single-board PC (Raspberry Pi 4 by Raspberry Pi Foundation) was used for real-time control. It received a 1kHz positional signal from the treadmill (also received by the Bpod), downsampling it to 25 Hz and then transferring it to a microcontroller (Arduino Nano by Arduino.cc) which controlled stepper motors to translate this into the synchronized motion of the plexiglass tube. If the mouse moved the treadmill faster than 4 cm / s then the downsampled signal was truncated to ensure that the tube followed smoothly. Note that the treadmill moved as fast as the mice needed; but if the treadmill moved at a high speed, the output was translated into a manageable speed for moving the lick spout, and the associated large tube that could contain a living rat.

### Rat stimulus & Visual stimuli

The position of the disk at the front of the tube was decided by the Bpod, and then updated by the single-board PC and the microcontroller (Arduino Nano by Arduino.cc), which controlled the stepper motors that rotated the disk. Finally, the single-board PC was used to generate the virtual visual environment tethered to the movement signal of the treadmill. The monitors used for displaying the streaming stimuli were also used as a go-cue at the beginning of the trial and an end-cue at the end. The streaming stimuli consisted of symbols that moved in synchrony with the treadmill. The screen flashed green at the start of a new trial and red at the end of a failed trial.

### Data acquisition

The treadmill sent out a 1 kHz signal encoding the current position. This signal was recorded on the state machine and stored to disk on a central workstation, along with the specific settings used in each trial. Two cameras were located below one of the screens used for the visual stimulus and set up such that they recorded a side view of the entire body of the mouse and its face, respectively, at 100Hz and sent this data to the central workstation. The workstation also provided centralized control over the single-board PC and state machine.

### Habituation and training

After surgical operation to implant a head post, mice were habituated to being handled and head-fixed (**Figure 1C, D**). Training on the treadmill began, once mice tolerated head fixation for ∼ 20 minutes. At this point, water intake for mice was monitored and restricted, but ensured to be at a level resulting in no more than 20% of weight loss per mouse compared to the weight before onset of water control. To ensure that all mice were available on the same experimental day when a rat was introduced, training continued for 3-4 weeks.

The initial training consisted of rewarding mice manually with Saccharose-sweetened water when they were head-fixed and they moved the treadmill in the correct / forward direction. Once mice moved the treadmill in the correct direction, training was automated. In the first days, there were no time constraints. Mice simply had to move at least 28.5 cm forward on the treadmill to receive the sweetened water reward, with no other limitation to speed or movement patterns. The total distance to the treadmill was 30 cm, but for safety reasons we automated the closing in of the spout on the last 1.5 cm. Five seconds after reward delivery, the lickspout / tube contraption automatically moved to its starting position, away from the mouse and the next trial began. The beginning of a new trial was indicated by a sound cue (played by a buzzer at 50Hz) and flashing of the virtual reality screens.

In the next phase of training, a time limit was introduced. Mice had 30 seconds to move the treadmill at least 28.5 cm and then another 2 seconds to lick the spout in front of them. Once mice learned to keep moving the treadmill to obtain a reward, the duration of the trial was shortened to 12 seconds. When the cohort of mice achieved >65% successful trials (with at least 30 trials per day) under these conditions for 3 days, the control and experimental data were collected.

### Live predator encounter

Once baseline control data had been collected on the next day (**Figure 1C-F**), a rat that had been habituated to handling and to the apparatus, was placed in the tube. The mouse was then head-fixed to the treadmill and experimental data were collected. The day after the encounter with the rat, a second day of baseline data was collected without the rat.

### Experimental paradigm with freely moving mice

One cohort of twelve male C57Bl6/J mice and a second cohort of twenty-four male C57Bl6/J mice were used in these experiments. Mice were ordered in equal numbers from the in-house breeding colony, Janvier, and Charles River. For the first cohort the 3 female Wistar-rats as in the simulated foraging task were used, and for the second cohort 2 additional female Wistar-rats were used. Female rats were used to minimize the possibility of a rat biting through the holder or injuring a mouse (Karli 1956). We chose to use animals from multiple sources to verify whether mice obtained from our in-house colony showed unusually low threat responses. For each session involving rats with freely-moving mice the experimenter randomly selected a rat to be used. To ensure that mice were naive to the stimulus in each of the paradigms, we ordered them by ascending complexity. We applied first the looming stimulus (artificial visual), then presented rat odor (natural olfactory), and finally a live rat (natural olfactory/visual/potentially tactile).

### Looming stimuli

To evaluate innate threat responses independent of the presence of a live predator, we applied the well-established looming stimulus paradigm. This paradigm is a standard for inducing fear in mice. The mice were solitary housed and habituated to changes in day-night-cycle for 5 days, then over the next four days, one mouse per source and day (resulting in 3 mice per day) was placed in a darkened room for an hour before experiments began. The purpose of the solitary housing of these mice was to ensure a minimum of unintended interaction between mice before each recording session. On the experimental day, mice were moved into a plexiglass arena (0.5 m x 0.29 m) and left to explore it for 10 minutes. The arena was cleaned with 15% Ethanol before each mouse was introduced. It contained a shelter, made of red plexiglass. When mice entered a zone encompassing the quarter of the arena floor most distant from the shelter (∼25 cm from the shelter edge), a looming stimulus (a shadow that expanded above the mouse) was presented. The looming stimulus consisted of 5 presentations of a small black dot (3°visual angle) expanding rapidly over 200ms to its full size (50°) and then remaining at this size for 250 ms. The stimulus was repeated at least 90 s after the last presentation. The sequence of looming stimuli was repeated 10-14 times for each mouse over 45 minutes. Video data was collected at 60 Hz. Trials ceased when a mouse stopped leaving the shelter for at least 15 minutes. Each stimulus presentation was triggered manually by a researcher supervising the experiment through a monitor located out of sight of the mouse, but within the same room as the setup. Consistency was ensured by defining a common criterion for activation of the stimulus. The trigger moment was defined as the moment when a mouse crossed a visible demarcation line (marking the distant quarter of the arena relative to the shelter) with all 4 paws.

### Test with Rat odor

The mice used in the looming experiments were then used to examine place preference in the presence or absence of rat odors. Over two days two mice from each source, i.e. in-house, Charles River, and Janvier, per day (6 mice per day) were moved into the same plexiglass arena as used in the looming stimulus test one at a time. The arena was split in half with a small connector between the two sides. Mice were left to explore both sides of the arena for 10 minutes. Then the mouse was removed, and a rat was brought into the arena, but could only explore one side of the arena. During this time the rat could move freely on that side of the arena. The rat was removed after 5 minutes, and the mouse was returned to the arena. If the rat urinated in the arena, the floor was wiped dry with a paper towel. Movement of the mouse in the two conditions was tracked offline with video acquired at 60Hz for the first cohort and 25Hz for the second cohort. The change in sampling rate was decided based on the data collected with the first cohort, as higher frame rates than 25Hz were not necessary for tracking the location of the mice. The arena was cleaned before each mouse was introduced into the arena.

### Assaying response to live rat

Finally, the same mice were placed in a modified version of the plexiglass arena (again 2 mice per group per day). The arena was divided in half by a 1mm thick metal mesh, with gaps that were just large enough for mice or rats to stick the tips of their noses through the mesh. Control data was acquired for five minutes, then mice were removed, and a rat was brought into the arena. Two minutes after introducing the rat, mice were introduced into the other side of the arena. Movement of the mouse was tracked offline with video acquired at 25-60Hz. The arena was cleaned before each mouse was introduced into the arena.

### Behavioral analysis

#### Simulated foraging with a live rat encounter predator encounter

Behavioral measures including overall performance, speed of movement and a variety of movement parameters were tracked using high speed video at 100 Hz, over the course of the 3 days. Analog traces from the treadmill encoder, and the movement sequences of the mice were captured on two Basler cameras, one aimed at the body, and one aimed at the face. Movement and posture of the mice was tracked using DeepLabCut (Mathis et al. 2018). Behavioral phenotypes that were available for analysis in the control condition and in the presence of the rat included: 1) Flight, apparent as a backward movement on the treadmill, and here detected specifically when a mouse moved at least 3cm backwards when directly in front of the tube. This movement pushed the tube holding the rat away from the mouse and showed up as negative deflection in the encoder output and led to a failure in licking. 2) Freezing, i.e. the mouse stops moving, which would be apparent in the speed traces and lead to a failure of the trial. This was detected here by either failing to complete a trial in time, or by failing to lick during the reward phase, when the mouse was directly in front of the tube. 3) Changes in mouse posture, hunching or stretching. 4) Changes in pupil size, detected through constriction or dilation of a circle fitted to the DeepLabCut-tracked edge of the pupil. 5) Changes in nose movement, which were also tracked with keypoints detected with DeepLabCut. Light conditions were verified to be generally stable and unrelated to pupil size in our recordings.

We performed the experiments in such a way that we were able to pair the behavioral phenotype of each mouse on the first recording day (before a rat was introduced) as a baseline to all further recordings of the same mouse.

#### Assaying threat response in freely moving mice

One of the three assays, the looming stimuli, was dependent on vision, two of them had olfactory components. In these experiments —looming, rat odor avoidance, and live rat exposure— the position of the mice was tracked using SLEAP (version 1.3.3). Three key body points were annotated along the midline of each mouse. For positional tracking, the pixel coordinates of the neck point were extracted and transformed into millimeter-based coordinates relative to the layout of each experimental arena. To reduce noise, positional data were clipped to the arena boundaries and smoothed using a sliding half-second window. Due to the side-view recording setup in the looming experiment (necessitated by an overhead screen), additional geometric corrections were applied. The X-coordinate was taken directly from the neck point, while the Y-coordinate was defined as the lowest Y-value among the three tracked points. An empirical downward shift of 20 pixels was then applied to the Y-coordinate. The adjusted coordinates were subsequently clipped to the nearest point within the arena polygon visible in each frame. After these corrections the pixel-based position is converted to millimeter-scale coordinates.

Because the perspective of the looming videos enforced a strong correlation between the X- and Y-position, only the X-position of the mouse relative to the shelter was used for further analysis. The absolute X-axis speed (relative to the shelter) was smoothed with an 11-frame window before being used to classify looms as either “fleeing” or “non-fleeing” events.

### Statistical analysis

#### Simulated foraging with live a rat

All statistical comparisons for eye position, pupil diameter, locomotor speed, posture, and facial movement were performed within the animal, comparing each mouse’s baseline session to its corresponding rat-exposure session. Time-resolved analyses were summarized into half-second bins, but statistical testing was conducted at the level of per-animal distributions rather than treating individual trials or bins as independent samples. Thus, each mouse served as its own control, avoiding pseudoreplication due to within-subject correlations. For time-resolved analyses, significance thresholds were adjusted for multiple comparisons across bins using a Bonferroni correction corresponding to the number of consecutive bins tested.

Here we first assayed mouse behavior for evidence of innate fear (i.e. freezing, fleeing) which would be evident in performance. This study was also designed to capture subtle changes in the behavior of mice (speed of movement as mice approach rats, success rate, pupil size, body elongation, nose movement) in the presence of rats. To assay these changes in behavior, the data related to each behavioral feature (pupil size, speed of movement, nose movement, etc.) were first filtered with a rolling z-score across a one second window. Values with an absolute score above three were removed. The time series data was then smoothed with a half second window by way of a rolling mean.

In the case of the video data (DeepLabCut output) it was also necessary to perform initial filtering steps, according to the detected likelihood that a feature was discovered in a frame, and with an initial z-score filtering across the x- and y-position of the features. In statistics where we considered the pupil-diameter and pupil position, we also filtered the eight detected markers around the pupil with a modified z-score (Iglewicz and Hoaglin 1993).

We compared the behavioral items pairwise per mouse, between the day of the baseline recording and the following days. To achieve this, we summarized half-second wide bins relative to the beginning of the trial or the delivery of reward of the time series and compared these between the two conditions. These data were plotted to show the mean and standard deviation per feature and condition. Binned data for sessions with and without the rat were compared, and the difference between the mean and standard deviation and the significance of this shift shown as a bar plot. Significance was assessed for each half second bin. Additionally, for each behavioral parameter (i.e. position, speed pupil size, eye position) differences between sessions with and without the rat had to be significantly different for 6 consecutive bins. In the reward phase, which lasted for a short duration, there had to be a significant difference over at least 3 consecutive half-second bins for the pupil x-position, pupil diameter, body length, and nose speed.

The continuous mean and standard deviation of the time series was calculated with the formulas 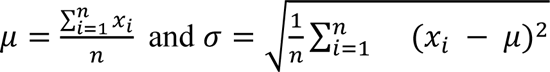. These values were then used to calculate the difference between both statistics. We used the following formulas for this: 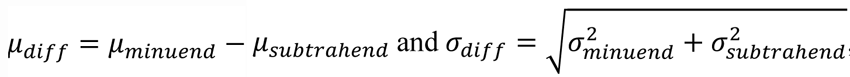, with the subtrahend always being the baseline condition and the minuend representing either the rat encounter condition or the measurements from the day after the simulated rat encounter. To determine the significance of the shifts, we used the Mann-Whitney U-Test. The original thresholds in the figures were *=0.05, **=0.01, ***=0.001, and n.s. = not significant, which were then applied with a Bonferroni-correction corresponding to the number of bins considered in each subplot.

For the summary data, multiple mice into one group, the means and standard deviations of their individual bins were combined, under the assumption of independence between mice, with the formulas 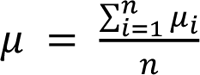 for the mean and 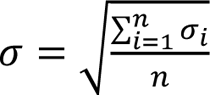 for the standard deviation of the combined bin. The p-values were averaged, weighted by the inverse of the variance of the individual bins they originated from. This happened according to the formula 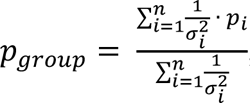, with n corresponding to the individuals in the group. Due to the high number of single samples per bin, the estimate of significance was limited to the ethologically relevant threshold of 10% of the 5^th^ to 95^th^ percentile span of each behavioral feature, across animals and conditions.

#### Data analysis for freely moving mice

All analyses were performed blind to the source of the mice. When shown, the unblinded mouse ID follows the format “C{Cohort #}{Source}{Mouse #},” where the source is indicated as C (Charles River), I (Internal), or J (Janvier). For example, the ID *C2J1* refers to the first mouse from Janvier in cohort 2. No significant behavioral differences were observed among mice from the three sources. Detailed categorizations by experiment, cohort, and source are presented in **Supplementary Figure 10**.

Mice were classified as “fleeing” or “non-fleeing” based on the median of all loom-wise ratios comparing the mean speed during the 5-second looming visual stimulus to the mean speed in the subsequent 5 seconds. A threshold ratio of 2 was chosen as a conservative criterion for identifying clear fleeing behavior.

The same threshold (a two-fold change) was used to conservatively identify mice that exhibited significant changes in location preference in the odor-based experiments. Specifically, this included avoidance or preference for the rat-odor side of the arena, as well as avoidance of the region adjacent to the mesh in the live rat exposure condition. Both behaviors relate to avoidance of rats as a defensive response. Mice falling below this threshold were classified as behaviorally unchanged.

### Data analysis Software

#### SLEAP

Two different SLEAP (Pereira et al. 2022) bottom-up models (version 1.3.3) were trained to track the location of the mice in the verification experiments. The side view model for videos recorded in the looming stimulus experiments was trained on 225 masked frames (to crop the moving looming stimulus and reduce the complexity of the video). The top view model for videos recorded in the rat odor and presence experiments was trained on 430 frames.

#### DeepLabCut

A series of DeepLabCut models (version 2.1; Mathis et al. 2018) were trained to track key features of mice for behavioral analysis in the simulated foraging task. Separate models were trained for each anatomical landmark: **Body key points** (nose, shoulder, tail base): trained on 220 labeled frames for one cycle of 1,030,000 iterations. **Nose key points** (nose ridge, nose base, nose tip): trained for two cycles of 1,030,000 iterations each. The first cycle used 220 labeled frames; the second used 337. **Eye key points** (eight equidistant points around the pupil, starting from the 12 o’clock position, plus the left and right corners of the eye): trained over three cycles of 1,030,000 iterations each. The training sets consisted of 260, 466, and 629 labeled frames, respectively. The networks were trained across animals and sessions, when performance was lower in a video we added more frames from that video to the training data.

## Results

### Simulated Foraging

This paradigm was designed to be easy for mice to perform but was also designed to minimize stress for both rats and mice. Once mice were habituated to head restraint and to obtaining reward by moving the treadmill (**Figure 1A,B,C**) they could perform 50-100 trials in a day, obtaining a total reward of 0.5 ml in a ∼30-45-minute session. When mice started a trial, the lickspout was at a starting position ∼30 cm from the mouse (**Video 1**, **Figure 1D**). A go cue, the flashing of the virtual reality monitors, and a sound cue initiated the trial. A successful trial was one in which mice moved the treadmill 28.5cm in 12 seconds, with the last 1.5 cm of movement controlled automatically within the code. Two to four seconds after reward delivery, depending on the start of licking, the lick spout and red tube over it were reset to their starting position.

Well trained, motivated mice started running on the tread mill immediately after licking the reward, even before the lick spout moved back to starting position. As mice improved their daily performance, adult female rats were habituated to being handled and to the apparatus. Once a cohort of mice were stable in their performance for 3 successive days, reaching the threshold criteria in their performance, the experimental data was collected over 3 consecutive days. On the first day control data was collected, on the second day the rat was introduced into the behavioral paradigm. The third day was a post-rat control session (**Figure 1E**). To maximize the potential for interaction between the head-fixed mouse and restrained rat, we used only one setting, open, on the disk that covered the mouth of the tube holding the rat. Note that in this setting mice could almost touch the rat on each trial and mice could see, smell and hear the rat moving in the tube as it came closer to the mouse. On each trial, with every mouse, rats could stick their nose out of the tube. When rats were in this position, their nose was effectively just above the lickspout for mice (**Video 1**). In the following plots the male mice are enumerated as mouse 1-4, and the female mice as mouse 5-7.

### Performance

Performance of each mouse was analyzed for three consecutive days: control / baseline day, day with the rat, and a post-rat day. Surprisingly, the success rate of most mice (5/7 mice) was unaffected (stayed above criterion for all three days) by the introduction of the rat (**Figure 1F**). Note that across the cohort of animals, performance was stable across days for the majority of animals. Five of seven mice maintained success rates above criterion (>65%) on the baseline day, during rat exposure, and on the post-rat day, with no systematic increase in trial failures or missed rewards when the rat was present (**Figure 1F**). In contrast, two mice showed a marked reduction in success rate during the rat session (>25% decrease relative to baseline), which persisted into the post-rat day, indicating a stable, individual-specific effect rather than transient trial-by-trial disruption. Importantly, the presence of the rat did not induce consistent hesitation at trial onset or prolonged latencies to initiate treadmill movement across the cohort; instead, changes in performance were confined to a small subset of animals. Next, we examined whether mice changed their behavior when the rat was introduced.

### Movement speed on treadmill

One measure of performance is the rate of success; another measure was the speed with which mice moved the treadmill (**Figure 2; Video 2**). Mice could run or walk slowly and consistently to cover 28.5 cm in 12 seconds. Most mice learned to move at a consistent speed of around 0.2 m / s. This speed was mostly uniform and on average it stayed constant for most of the trial (**Figure 2B**), but the speed decreased abruptly as mice stopped moving to lick the reward spout (**Figure 2B**).

**Figure 2.**
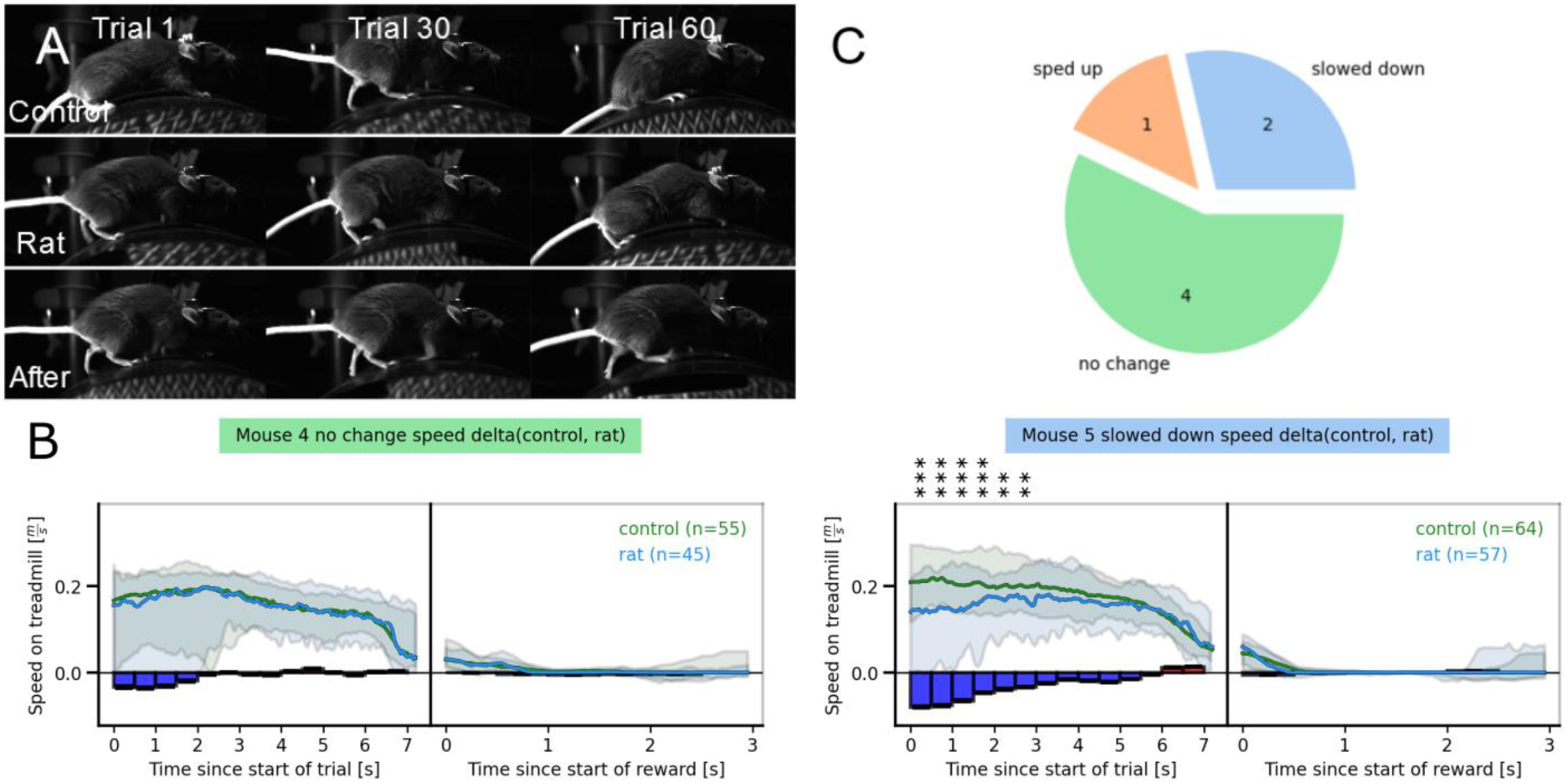
Movement speed in the presence of rat. **A**) Representative images of mouse position from three sessions. The images show the position of a mouse at 3 different times on three different sessions. These kinds of images were used to assess whether the head-fixation height or position of the mouse were stable from day to day. Plots for each of the seven mice are in **Supplement-figure 8**. **B**) Speed of movement on the treadmill. The median and confidence interval of the speed of movement in baseline, control day, recordings (black trace) and the session with the rat present (blue) show the two outcomes of having the rat hovering over the lick spout. The left plot (trial phase) shows no change in velocity throughout the trial. In another mouse there was a momentary reduction in speed at the beginning of the trials (right panel, trial phase). When the mice were at the lick spout, they stopped moving (see reward phase). Significance was assessed with a Mann-Whitney U-test (*: p<0.05, **: p<0.01, ***: p<0.001). See methods for additional filtering applied to significant effects. **C**) Pie chart grouping mouse behavior. Mice could be divided into two main groups, one that showed significant changes in speed and another that showed no change in speed. One mouse significantly increased its speed during the approach to the tube with the rat, 4 mice showed no significant change of speed while approaching the rat, and 2 slowed down significantly.

One issue that arose in monitoring and comparing the behavior of mice from day to day, was whether the position of the mouse on the treadmill changed across days. It was possible that mice were positioned at slightly different heights relative to the treadmill, and this effectively changed the speed with which mice moved (**Figure 2B**). To examine this, we selected frames from different time points during each session and examined them for any obvious differences in height of the mouse relative to the treadmill (**Figure 2A; Supplementary Figure 9)**. There were no significant differences in the positioning of mice from day to day.

Next, we compared the speed of movement from control sessions (**Figure 2B**, green traces) and sessions when the rat was present for individual mice. These data show that while the majority of mice show no consistent significant changes in their movement speed in presence of the potential predator (**Figure 2B**, blue traces), three out of seven mice showed significant (p<0.05, binwise Mann-Whitney U (MWU) test, Bonferroni Correction) and consistent changes in their movement speed throughout the session (**Figure 2C**). Their average speed was significantly different at the onset of the trials when the rat was present. The two mice that decreased their movement speed in the presence of the rat were slow on the treadmill from the beginning of the trial. One mouse ran significantly faster in the presence of the rat. In the remaining mice, there were no consistent changes in average speed (**Supplementary Figure 3, 4**). Thus, while the presence of the rat altered locomotor speed in a subset of mice, these effects were directionally heterogeneous (slowing in two mice, speeding in one) and did not reflect a uniform suppression of movement or task engagement.

### Eye movement and pupil diameter

To assess whether mice attend to the presence of the rat, we used DeeplabCut to track eye movement and pupil diameter on the control day and on the day that the rat was introduced into the tube (**Figure 3, Video 3**). We plotted the average position of the pupil in the horizontal axis relative to the two corners of the eyes, over the course of the control session and in the presence of the rat (**Figure 3A-C**). In five out of seven mice, there was a significant (p< 0.05, binwise MWU, Bonferroni Correction) change in the horizontal position of the pupil in the presence of the rat (**Figure 3B**, green traces); in the remaining 2 there were no consistent changes in eye position. On average, the position of the eyes of five mice was significantly different (p < 0.05, binwise Mann Whitney U Test, Bonferroni correction) during their reward phase compared to the control days (**Supplementary Figure 5**). When the rat was present in the tube and the mice were directly in front of the tube, on average, 3 mice positioned their eyes more in the direction of the rat (looked right) and 2 looked away from the rat. These effects were significant and were evident throughout the trial: across mice that showed significant effects, horizontal eye position shifted by an average of ∼1-3% of the half width of the eye relative to baseline. Two mice showed no significant change in their eye position (**Figure 3C**, right panels).

**Figure 3.**
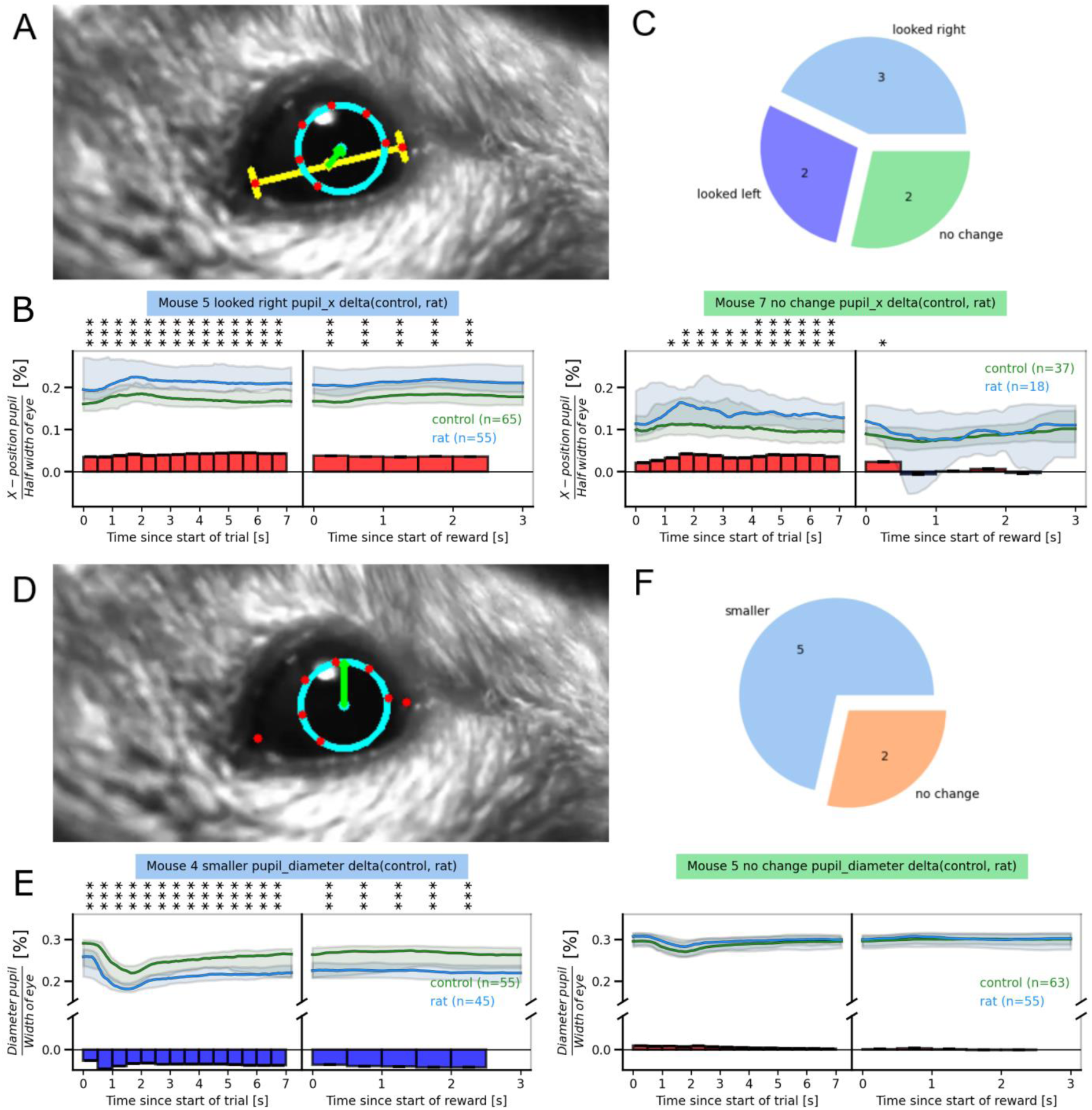
Pupil movement and pupil size in the presence of the rat. A) Schematic showing the analysis of horizontal pupil position. The x-position was defined relative to the center of the pupil and the corners of the eyes. Movement of the eyes could be nasally oriented (toward the rat) or oriented to the back of the head away from the rat. **B)** Plots for a mouse that positioned its eyes toward the lick tube (nasally) as it moved toward the lick spout (left plot) and positioned its eyes to look forward toward the rat, when the mouse was stopped at lick spout (right plot). The second set of plots on the right show eye position for a mouse which showed no significant change in x-position while in front of the rat. **C**) Pie chart grouping mice. Five mice positioned their eyes significantly differently in the presence of the rat (n=5). Two mice showed no change. Three mice looked toward the tube holding the rat more often, and two looked away from the rat. **D**) Schematic of the pupil diameter analysis. The diameter was calculated based on points labeled by DeepLabCut. **E**) Example plots from two mice, one showing a significant reduction in pupil diameter in presence of the rat (black trace is control session, blue trace was with rat present) the other one showing no effect. **F**) Pie chart grouping mice. Mice could be divided into two groups, mice that showed no change in pupil diameter and a second group of five mice that had a consistent reduction in pupil diameter in the presence of the rat.

Next, we examined pupil size (**Figure 3D-F; Video 4**). Pupil size changes with changes in lighting, or changes in the parasympathetic or sympathetic system. To establish that light around the face and head of the mouse remained consistent, we measured brightness and light intensity around the eye and at another point on the head. There was no effect on the luminance / light levels around the eyes or head when the rat was introduced. Our data suggest that across mice that showed a significant reduction, pupil diameter decreased by ∼10-13% during successful approaches, with these changes remaining stable across trials within a session (Fig. 3; Supplementary Figs. 5–6).

But in the presence of the rat there were significant (p<0.05, binwise MWU, Bonferroni correction) changes in pupil diameter in five out of seven mice (**Figure 3D, Supplementary Figure 6**), especially when mice were close to the rat (Blue traces, **Figure 3E**). In these mice, the pupil diameter was consistently smaller throughout the course of the trial and in four out of five of these mice the change in pupil size persisted into the next session. The other 2 mice showed no significant changes in their pupil diameter. Note that the reported significance values reflect within-animal comparisons between baseline and rat sessions, with each mouse treated as an independent observational unit. Taken together, these data show that mice detect and respond to the predator, with pupil dynamics consistent with an altered attentional or autonomic state.

### Posture and facial movements

To assess whether any other aspects of mouse posture or facial movement were affected by the presence of the rat, we measured mouse posture, estimated by body-length, and facial movements (**Figure 4A-C; Video 5, Supplementary Figure 7, 8**). In three out of seven mice there was a significant change in the pose when the mouse was confronted by the rat (blue). Additionally, in 2 other mice there was a significant increase in nasal movement speed in the presence of the rat (**Figure 4D-F; Video 6**).

**Figure 4.**
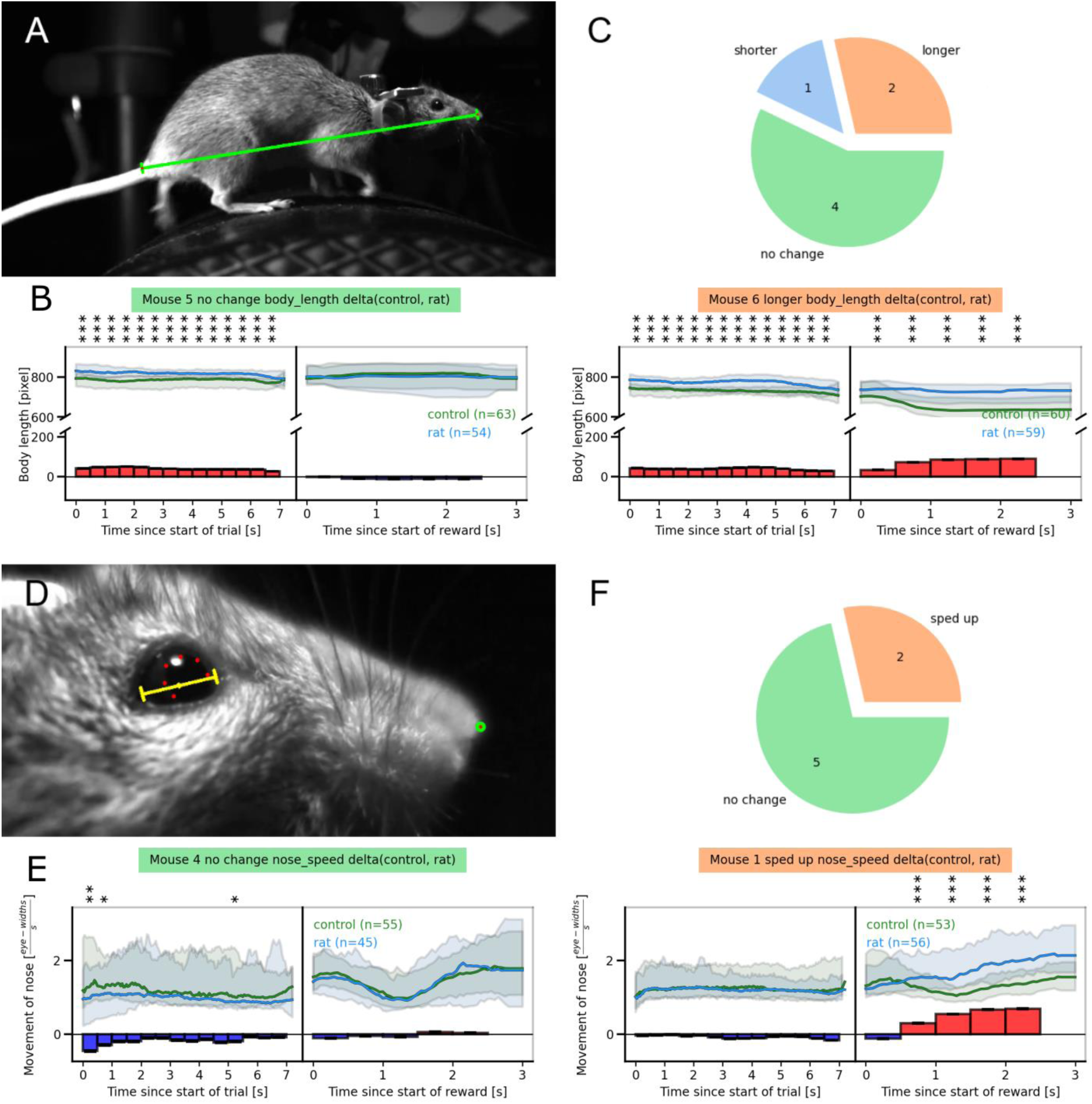
Changes in posture and facial movement in the presence of a rat. **A**) Schematic showing analysis of posture using a tail to nose distance. **B**) Body-length diagrams for two mice. The mouse on the left showed no significant change in posture when in front of the rat, the one on the right was significantly more hunched up in the presence of the rat. **C**) Pie chart grouping mice. Mice could be divided into three groups: 4 mice showed no change in body length in the presence of the rat, and 2 mice were more hunched up at the lick tube in the presence of the rat, and one elongated its body at the lick spout in the presence of the rat. **D**) Schematic showing the analysis of facial movement / nose speed. The nose position was detected with DeepLabCut and normalized to the distance between the corners of the eye, to account for mouse size and position. **E**) Facial movement. The left plots show data from a mouse that did not significantly change the movement speed of its nose even when it was directly in front of the rat. The right-side plots show movement data from a mouse that increased the movement speed of its nose significantly, during the reward phase. **F**) Pie chart grouping mice. Most mice showed no significant change in nose speed even when they were directly in front of the rat.

Taken together, these results show that although head-fixed mice do not express canonical escape behaviors, the presence of a live rat reliably alters multiple behavioral and physiological variables, including locomotor dynamics, eye position, pupil diameter, posture, and facial movements (**Supplementary Figure 10**). These changes indicate that mice detect and respond to the predator, but express this response through a constrained behavioral repertoire rather than through overt flight or freezing. Note that although both male (n = 4) and female (n = 3) mice were included in the head-fixed experiments, no consistent sex-specific differences were observed in task performance, locomotor speed, eye movements, pupil responses, or posture; given the small sample size, sex was not treated as an independent factor in statistical analyses.

Next, we considered whether head fixation, thirst, and the need to forage affected threat perception or the mouse’s behavioral response to the threat. At the same time, we also assessed whether the mice in our colony were inherently less fearful. To address these issues, we used naive freely moving mice, from our colony and from two other sources.

### Effect of looming stimuli

In these experiments, we used 36 naive mice, 12 mice were from our animal facility, 12 newly acquired from Charles River, and 12 from Janvier. Mice obtained from our inhouse facility showed no differences in behavior compared to mice obtained from external sources.

Mice were placed in an arena (**Figure 5A, B**), and when they entered a particular location in the arena, the looming protocol was initiated. A dark shadow emerged, and grew larger, hovered over the mouse, simulating a bird of prey swooping over the mouse. For each mouse, the looming stimulus protocol was repeated 14 times. Post-hoc video analysis showed that none of the mice froze in response to the looming stimulus; seven out of 36 mice ran into the shelter (**Figure 5C, D**). The seven mice that ran into the shelter moved rapidly toward the shelter during the looming stimulus (**Figure 5E, F**), the remaining 29 mice did not show any significant, consistent movement toward the shelter. Overall, these results suggest that looming stimuli can reliably evoke fleeing in mice, but this effect is only observed in ∼19% (7 out of 36, Wilson 95% confidence interval of ∼10–35%) of the mice.

**Figure 5.**
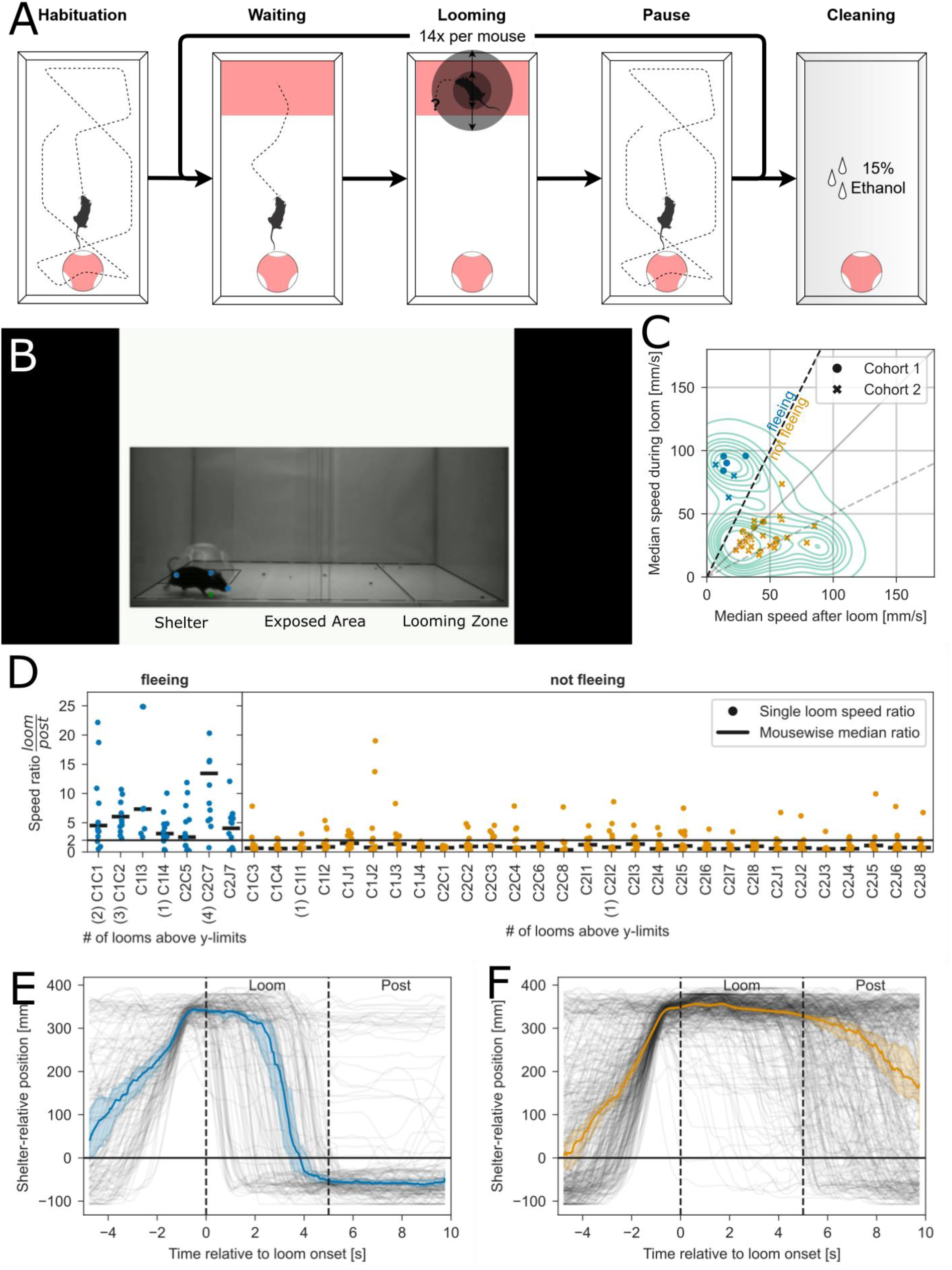
Effect of looming stimuli on the behavior of freely moving mice. **A)** Experimental protocol for the looming stimulus paradigm. **B)** Representative annotated video frame. Black bars denote frame boundaries, and the white area marks the masked region used to isolate the tracking area. The transformed arena outline is overlaid. **C)** Scatterplot showing the median shelter-relative speed for each mouse across all loom presentations. The behavior classification threshold (speed ratio = 2) is indicated by a dashed black line. Individual cohorts are marked with distinct symbols, and behavioral groups (“fleeing” vs. “non-fleeing”) are color-coded. The global distribution of all average speeds is shown as a background contour plot. Seven of 36 mice met the criterion for consistent flight behavior (see Methods). We avoided defining a “freezing” group based on these parameters, as the trial-wise speed distribution did not show a clean separation as it did for the “fleeing” animals. **D)** Strip plot of average speed ratios per mouse. The median ratio for each animal is shown as a black horizontal bar. Mice were classified as “fleeing” if their median ratio exceeded 2. Numbers in brackets next to mouse IDs indicate how many individual looms exceeded the plot’s y-axis limit. Mouse *C1I3* completed only seven looms, having remained in the shelter after the seventh presentation. **E)** Time-aligned traces of shelter-relative X-position for all loom events from mice in the “fleeing” group. The median and 95% confidence interval are shown as a bold line and surrounding shaded region, respectively. **F)** Same as in E, but for loom events from mice in the “non-fleeing” group. Vector illustrations of the rat and mouse in this and subsequent figures were adapted from SciDraw (SciDraw, 2020a; Costa, 2020).

### Avoidance of rat odors or rats

Next, we examined whether freely moving mice (n = 36) reacted to rat odors (Figure 6A-C). Mice were placed in an arena with a partition down the middle, a partition that mice could and did traverse. Five minutes after putting mice in the arena, they were removed from the arena. A rat was introduced on one side of the partition, a partition that rats could not traverse. Before mice were returned to the arena, the rat was removed. Post-hoc analysis of the amount of time spent in each portion of the arena, revealed that most mice either preferred the side that had contained the rat (6 mice out of 36), or showed no clear preference for either side of the arena (29 mice out of 36) (Figure 6C).

**Figure 6.**
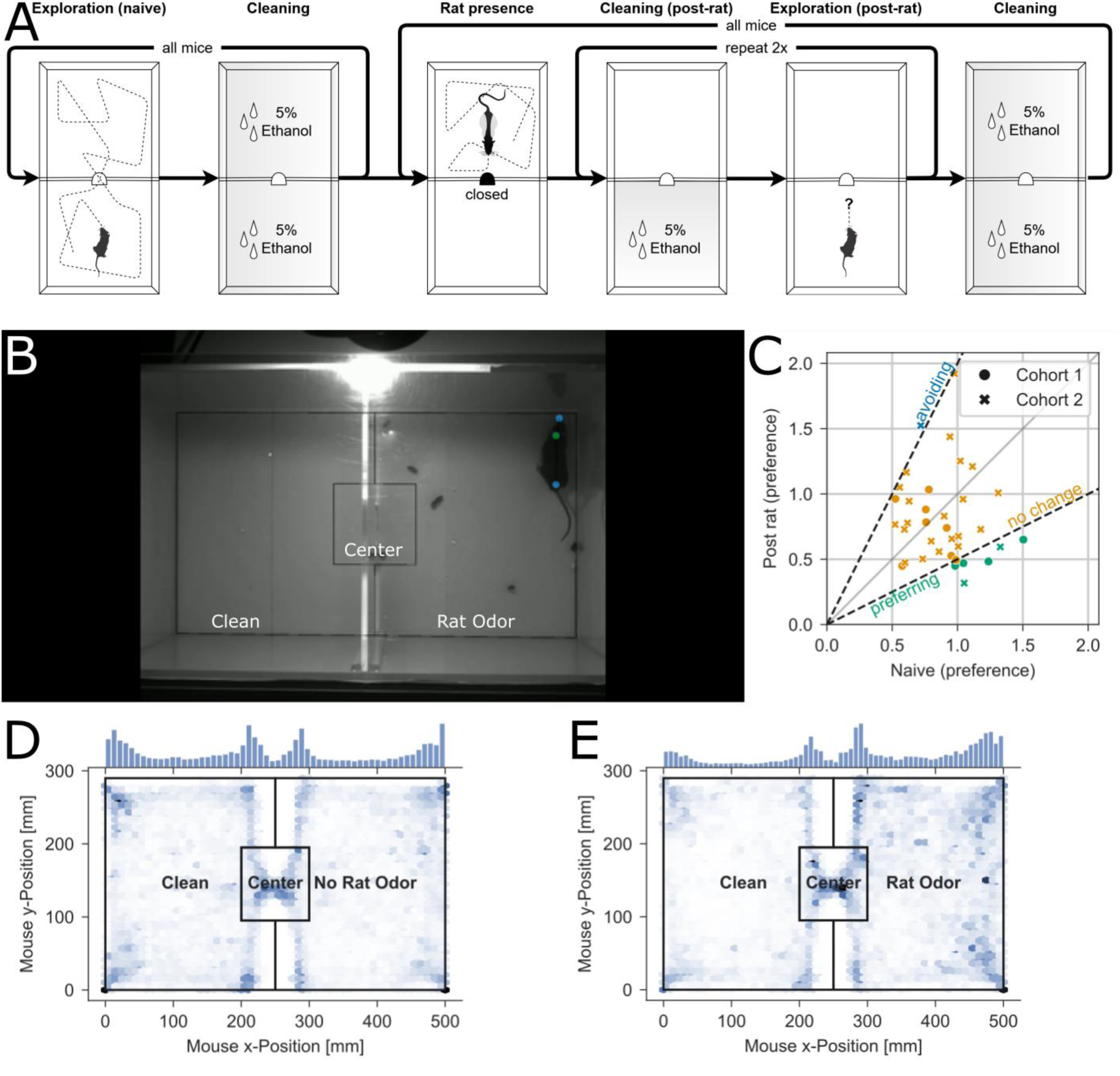
Avoidance of rat odors. **A)** Protocol of rat odor avoidance experiment. **B)** Same as Fig. 5B, but for the rat odor avoidance experiment. **C)** Scatterplot mapping mouse-wise preference for the clean area before and after the rat was introduced to the “rat odor” area. The behavior class criteria are shown as dark black dashed lines, while the 2 cohorts of animals used in the experiments are marked with different symbols. The resulting behavior groups are shown with three different colors (avoiding the rat odor in blue, which did not occur); and no change in preference (gold) and preferring the rat odor (green). Six out of 36 mice preferred the area with rat odor after the rat was introduced. **D)** Overall distribution of positions of mice in the arena, belonging to the group of mice “preferring” the rat odor, before the rat was introduced. **E)** Same as D, but for the position distribution after the rat was introduced.

We used the same arena but closed off the wire mesh partition that separated the arena into two halves (**Figure 7A, B**). Once again naive mice were placed in the arena but were allowed to explore only one half of the arena, then they were removed, and a rat was placed on the right side of the arena. Mice were then returned to the arena. In these experiments, half the mice (n=18) avoided being close to the wire mesh that separated the rat from the mice (**Figure 7C-E**). Of the other half (n=18) fifteen mice showed no preference for one side and three mice preferred staying close to the mesh. These data indicate that exposure to a live rat does not elicit a uniform, automatic defensive response across mice. Instead, freely moving mice exhibit substantial inter-individual variability, including avoidance, investigatory behavior, and neutral exploration, consistent with context-dependent threat evaluation rather than a fixed innate response.

**Figure 7.**
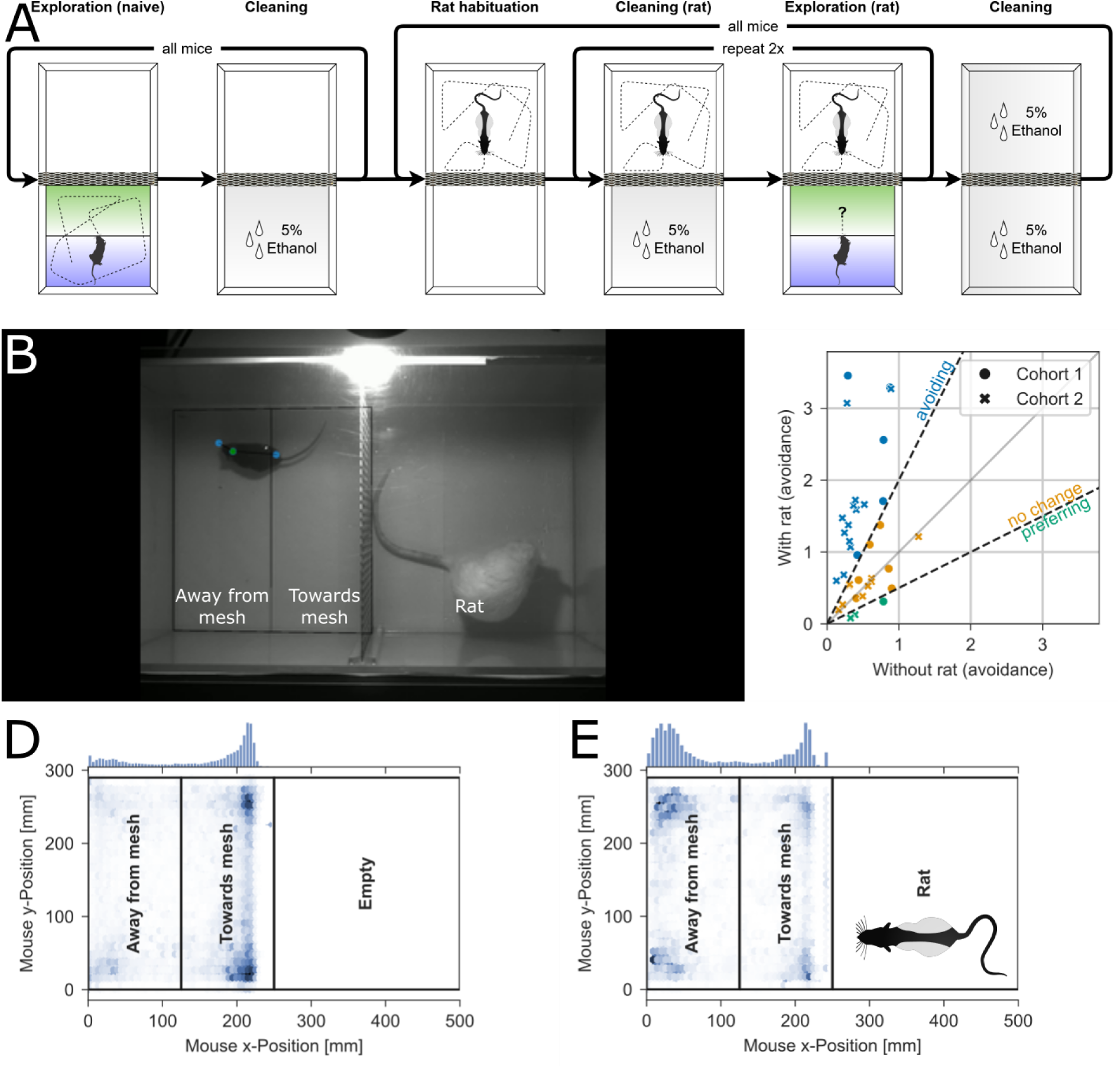
Avoidance behavior in the presence of a live rat. **A)** Protocol of rat presence experiment. **B)** Same as Fig. 5B, but for the rat presence experiment. **C)** Same as Fig. 6B but for the ratio of avoidance of the area close to the mesh without and with the rat present. All three possible behavioral groups were observed (avoiding in blue, preferring in green and no change in yellow). Eighteen out of 36 mice avoided the mesh when the rat was present, while 3 mice preferred to explore the mesh when the rat was present. **D)** Same as Fig. 6D but applied to mice that avoided the mesh once the rat was present, showing positions before the rat was present. **E)** Same as D but showing the overall position distribution while the rat was present.

## Discussion and conclusion

Defensive behaviors such as freezing, fleeing, or fighting are often treated as automatic responses to predator threat. Our results challenge this assumption by showing that predator exposure does not necessarily elicit an invariant defensive repertoire in mice, even under ethologically relevant conditions. Across both head-fixed and freely moving paradigms, mice frequently did not express stereotyped defensive actions yet showed clear evidence of threat engagement through changes in locomotion, gaze direction, pupil diameter, and posture. Together, these observations suggest that defensive behavior can be expressed flexibly and is not obligatorily triggered by predator cues.

In the head-fixed simulated foraging task, mice were physically unable to flee, and freezing would have resulted in trial failure and loss of reward. Under these constraints, most mice continued to perform the task despite close proximity to a live rat. Importantly, continued locomotion did not imply an absence of threat detection. Instead, individual mice exhibited reliable changes in multiple behavioral and physiological variables, including altered treadmill speed, systematic shifts in eye position, sustained pupil constriction, and changes in posture and facial movements. These measures indicate that mice detected and responded to the predator, but their response was restricted by head fixation and competing motivational demands of reward acquisition versus threat avoidance. Although the number of head-fixed animals was necessarily limited, each mouse contributed a large number of trials per condition, enabling sensitive within-animal comparisons, while constraining population-level generalizations.

This interpretation helps reconcile our findings with previous work. Vollmer et al. (2022) reported robust suppression of reward-seeking in head-fixed mice exposed to predator-related odors (e.g., TMT) under conditions in which animals were stationary, had ad libitum access to food and water, and reward-seeking was operationalized as discrete lever presses. In contrast, mice in our paradigm were water restricted, actively controlled their approach toward the stimulus, and were exposed to a live predator rather than an olfactory cue alone. Reward acquisition additionally required sustained locomotion followed by licking, rather than a single action. These differences in motivational context, threat modality, and action requirements are likely to strongly influence how defensive states interact with reward-seeking behavior, indicating that suppression of reward seeking under threat is context dependent rather than obligatory.

The freely moving experiments in this study provide an important complementary reference point. Using established ethological assays—including looming stimuli, predator odor, and live predator presence—we observed pronounced inter-individual variability and strong context dependence, with defensive responses expressed in only a subset of animals. These experiments therefore serve as a positive control, indicating that the absence of uniform freezing or escape in the head-fixed paradigm does not reflect insensitivity to threat. To assess whether head fixation, motivational state, or colony-specific factors contributed to this pattern (Burnett et al. 2016; Verma et al. 2016), we tested freely moving, well-fed mice drawn from multiple sources. Even under these conditions, looming stimuli elicited escape behavior in only a minority of animals (Lenzi et al. 2022), predator odor did not produce consistent avoidance (Banik and Anand 2011), and exposure to a live rat resulted in avoidance in approximately half of the mice (Karli 1956; **Supplementary Figure 11**). Together, these findings mirror the heterogeneity observed in the head-fixed paradigm and argue against a simple, hardwired mapping between predator cues and defensive motor programs.

The observed pupil constriction in the presence of a predator was unexpected, as threat is often associated with dilation. However, pupil size reflects a balance between sympathetic and parasympathetic activity and is sensitive to attentional state. In this context, constriction may reflect heightened visual focus or vigilance during close-range threat monitoring rather than reduced arousal. This interpretation is consistent with the idea that mice adopt alternative physiological strategies when classical defensive actions are unavailable or maladaptive.

As with many behavioral paradigms, responses to predator-related stimuli are shaped by prior experience, social history, and internal state (Yilmaz and Meister 2013; Lenzi et al. 2022; Wang 2020; Qi et al. 2025). Although we followed established protocols closely (Yilmaz and Meister 2013; Banik and Anand 2011; Karli 1956), differences in housing, handling, and individual experience are known to influence defensive expression. In earlier looming paradigms, high rates of freezing or escape were reported, but these effects depended strongly on stimulus parameters and arena structure, such as the availability of shelter. Defensive responses also vary with threat history and repeated exposure, including rapid habituation or suppression across trials (Yilmaz and Meister 2013; Lenzi et al. 2022; Qi et al. 2025). Together with prior work showing experience- and history-dependent modulation of looming responses, our results indicate that variability in defensive expression reflects flexible strategy selection rather than experimental noise.

### Caveats and issues to consider

In the present study, mice were socially isolated for several days before testing and were drawn from multiple sources, yet defensive responses remained limited and heterogeneous across a relatively large cohort (n = 36). Rather than treating this variability as experimental noise, our results suggest it is a fundamental feature of defensive behavior, reflecting flexible strategy selection shaped by context and experience rather than a reflexive, invariant output.

Because the same cohort of mice was tested across paradigms, experience-dependent carryover effects cannot be fully excluded. However, if such effects were present, prior exposure to predator-related stimuli would be expected to enhance, rather than suppress, subsequent defensive responses. The limited and heterogeneous defensive behaviors observed across tasks therefore argue against sensitization-driven inflation of fear responses and support the conclusion that defensive expression is strongly context dependent.

Note that in the head-fixed paradigm, the identity of the rat placed in the tube was rotated across sessions to minimize systematic effects of individual rats. In the freely moving experiments, rats were selected at random from a small set of animals (three rats in cohort 1 and two rats in cohort 2). Differences in rat behavior could, in principle, influence the salience of the predator stimulus and thus represent a potential source of variability. However, we observed no qualitative or quantitative differences in mouse behavioral outcomes between the two freely moving cohorts (**Figures 5–7C**), despite the use of different rats. This consistency argues against rat identity being a major determinant of the observed patterns of defensive behavior and suggests that the reported effects reflect mouse-specific and context-dependent factors rather than systematic differences in predator behavior.

It is possible that some aspects of how we habituate, or house animals or how animal caretakers look after mice reduce fear and stress or reduce the behavioral expression of fear in mice (Gouveia and Hurst 2017; Furlong et al. 2016; Kallnik et al. 2007). It is also possible that just as in rats, the sexual identity of mice affects how they respond in our behavioral paradigms (Gruene et al. 2015).

A key limitation of head-fixed paradigms is the restriction of overt defensive actions such as escape or freezing. Even though mice could have stopped walking forward, which would have the effect of keeping the rat and the reward at distance, or could have walked backward to push the rat away, they did not. Rather than viewing this as a failure of the model, we emphasize that this constraint enables a complementary question: whether threat detection and defensive state expression necessarily require these actions. Our findings suggest they do not. Even when classical motor outputs are unavailable or maladaptive, mice engage alternative behavioral and physiological strategies consistent with active threat assessment and coping.

Our study challenges simplified views of innate fear as a fixed behavioral program and highlights the importance of considering how environment and task demands shape the expression of defensive states.

## Table of abbreviations

MWU: Mann-Whitney U Test

## Acknowledgements

We thank the Charité Workshop for technical assistance, especially Alexander Schill, Jan-Erik Ode and Daniel Deblitz. We also want to thank Melissa Long and the team of the ABPF for their help with animal handling and room planning. Finally, we thank Laura Schwarz of the Sainsbury Wellcome Centre and members of the Larkum lab for useful discussions about earlier versions of this manuscript.

## Data availability statement

The original contributions presented in the study are publicly available. This data can be found here: http://doi.org/10.5281/zenodo.15264865. Extended data is available on request.

## Code availability statement

The code used to generate the data shown in this study is publicly available. It can be found here: https://github.com/Marti-Ritter/contextual-modulation-and-blunted-defensive-responses-to-predators.

## Ethics statement

All experiments were conducted in accordance with the guidelines of animal welfare of the Charité Universitätsmedizin Berlin and the local authorities, the ‘Landesamt für Gesundheit und Soziales’.

## Author Contributions

|  | Conceptualization | Data curation | Formal analysis | Funding acquisition | Investigation | Methodology | Project administration | Resources | Software | Supervision | Validation | Visualization | Writing – original draft | Writing – review & editing |
| --- | --- | --- | --- | --- | --- | --- | --- | --- | --- | --- | --- | --- | --- | --- |
| MR | x | x | x |  | x | x | x |  | x | x | x | x | x | x |
| LC | x | x |  |  | x | x |  |  |  |  |  |  |  | x |
| LS | x | x |  |  | x | x |  |  |  |  |  |  |  | x |
| AH | x | x |  |  | x | x |  |  |  | x |  |  |  | x |
| SÖ |  | x |  |  | x |  |  |  |  |  | x |  |  | x |
| RB | x |  |  |  |  | x |  |  | x |  |  |  |  | x |
| AV | x | x |  |  | x | x |  | x |  |  |  |  |  | x |
| DS | x |  |  | x |  |  |  |  |  |  |  |  |  | x |
| PP | x |  |  | x |  |  |  |  |  |  |  |  |  | x |
| ML | x |  |  | x |  | x |  | x |  |  |  |  |  | x |
| RS | x |  |  | x |  | x | x | x |  | x |  |  | x | x |

## Generative AI statement

The authors declare that no Generative AI was used in the creation of this manuscript.

