## Supplementary Figures and Captions for "Context-dependent reshaping of defensive responses to predators in head-fixed and freely moving mice"

No. of Figures: 11.

No. of Videos: 6

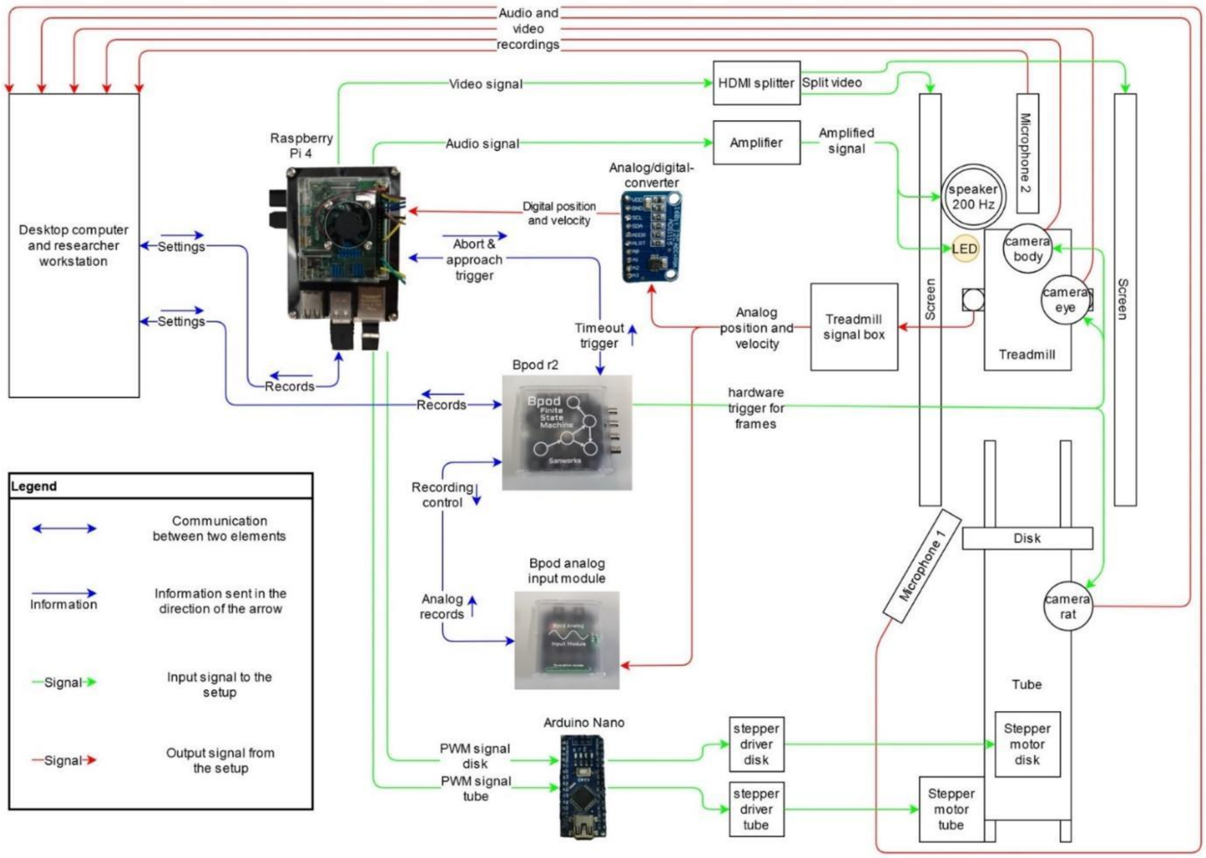


**Supplementary Figure 1.** *Connections in the simulated foraging setup.* The simulated foraging paradigm (see Methods) was run in the shown setup. Signal and data flows are marked in color.

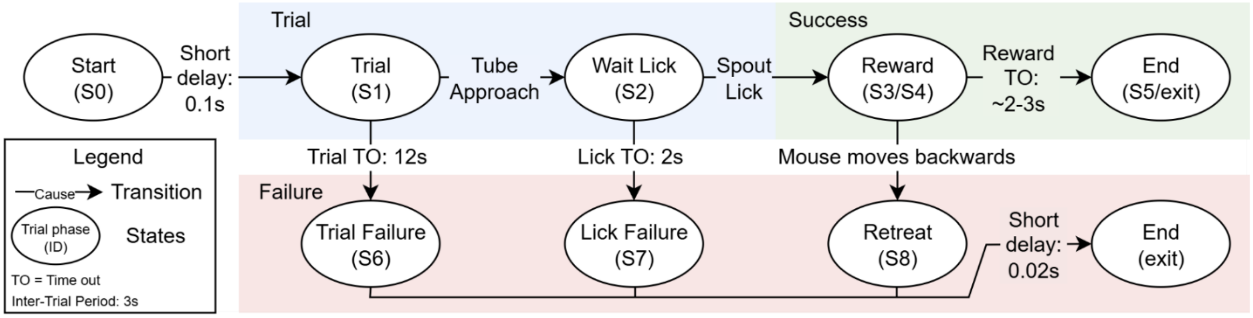


**Supplementary Figure 2: State machine controlling the simulated foraging task.**

Each trial begins in State 1 (S1). If the mouse traverses the required distance of 28.5 cm on the treadmill within 12 s, the lick spout is reached (S2). At this point licking the spout will trigger reward delivery via solenoid activation (S3–S4). Following reward delivery, a timeout (TO) is used to allow the mouse to lick the remaining reward and the trial terminates (S5). After a 3s inter-trial interval, the next trial begins. If the mouse fails to traverse the 28.5 cm within 12 s, S1 times out and is classified as unsuccessful (S6). If the mouse reaches the lick spout but fails to lick before S2 times out, the trial enters a separate failure state (S7). Backward movement of the treadmill during the reward states (S3/S4) also transitions the state machine to a distinct failure state (S8).

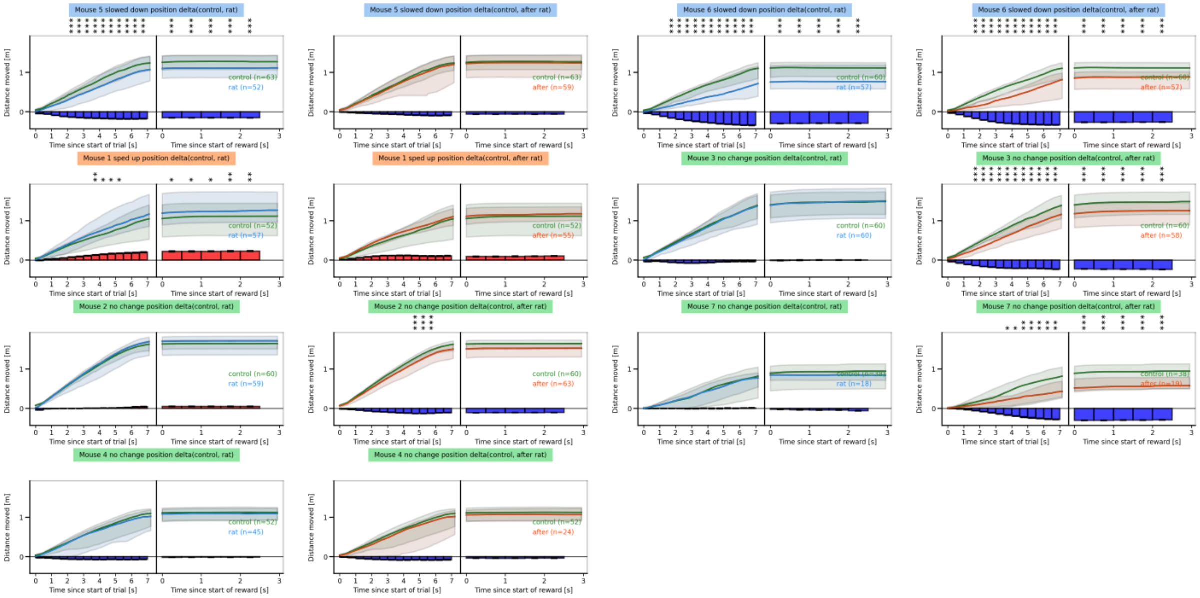


**Supplementary Figure 3.** *Analysis of position over time for individual mice.* Position analysis for all 7 mice in the simulated foraging paradigm. Same visualization as in Figures 2-4.

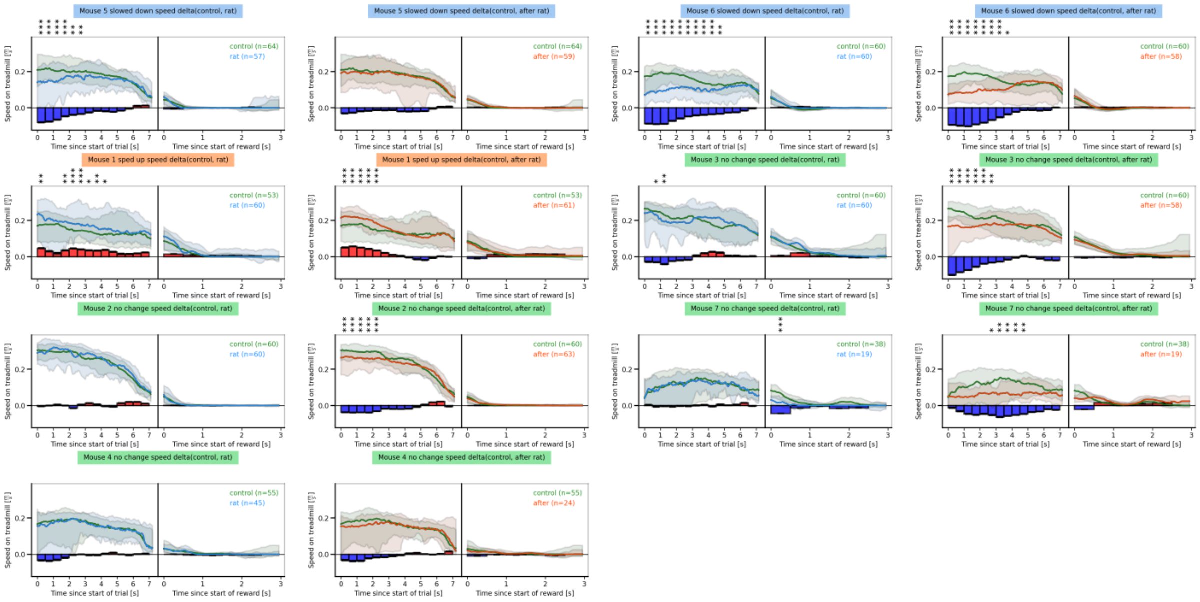


**Supplementary Figure 4.** *Analysis of speed over time for individual mice.* Velocity analysis for all 7 mice in the simulated foraging paradigm. Same visualization as in Figures 2-4.

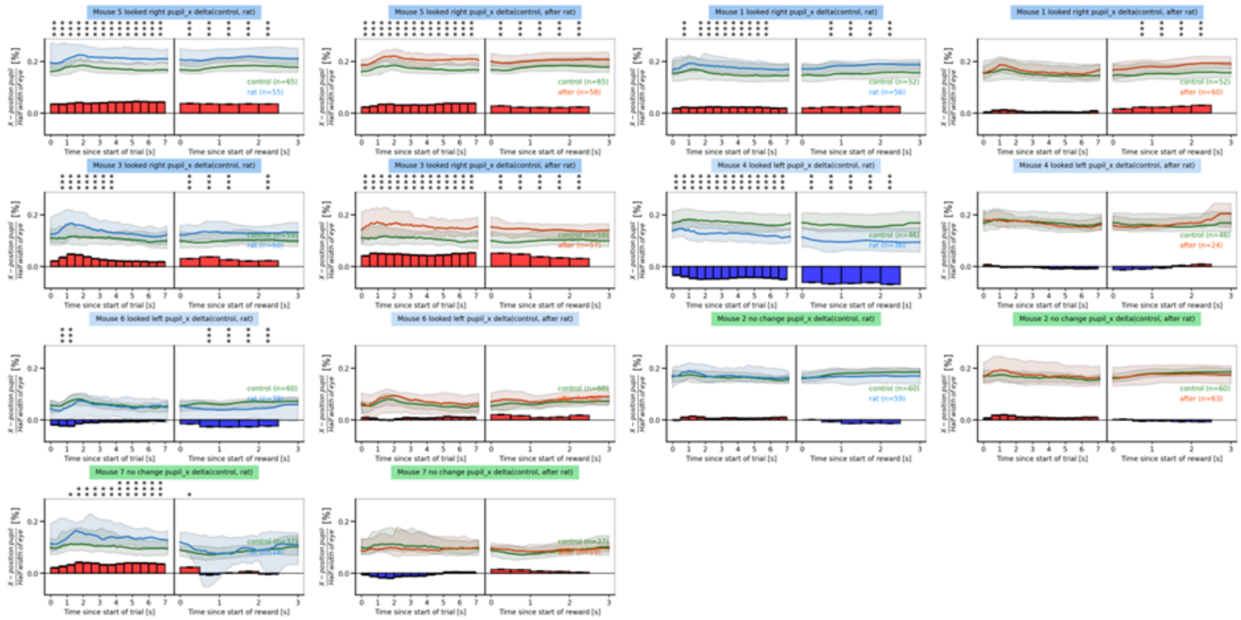

**Supplementary Figure 5.** *Analysis of horizontal pupil position over time for individual mice.* Pupil position analysis for all 7 mice in the simulated foraging paradigm. Same visualization as in Figures 2-4.

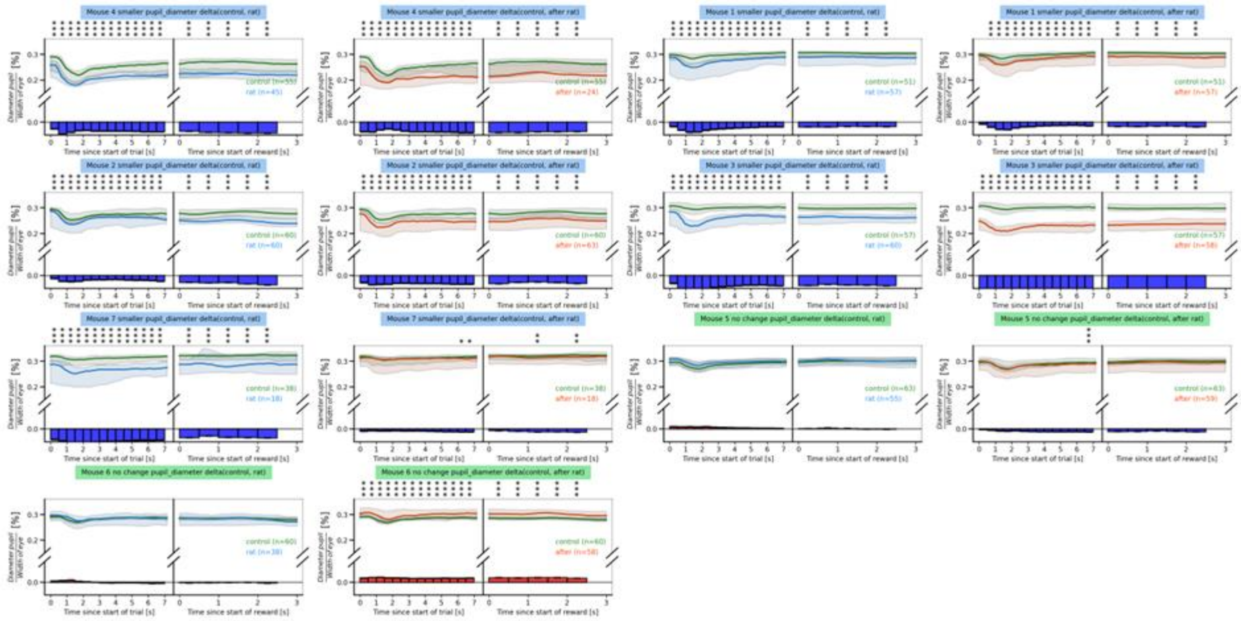


**Supplementary Figure 6.** *Analysis of pupil diameter over time for individual mice.* Pupil diameter analysis for all 7 mice in the simulated foraging paradigm. Same visualization as in Figures 2-4.

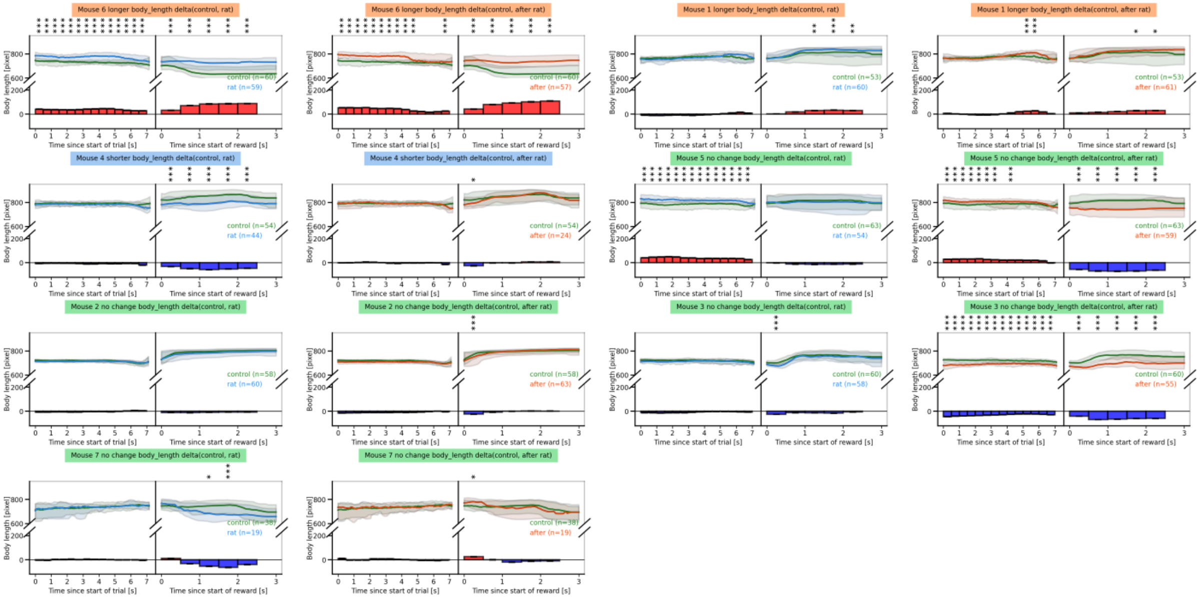


**Supplementary Figure 7.** *Analysis of body length over time for individual mice.* Body length analysis for all 7 mice in the simulated foraging paradigm. Same visualization as in Figures 2-4.

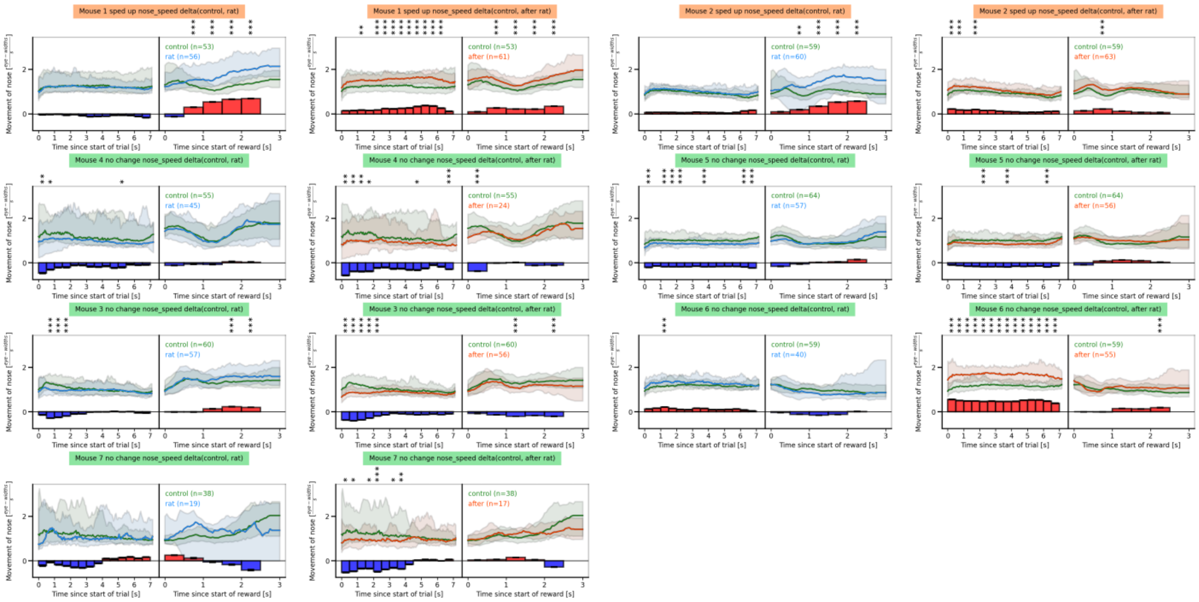

**Supplementary Figure 8.** *Analysis of nose speed over time for individual mice.* Nose speed analysis for all 7 mice in the simulated foraging paradigm. Same visualization as in Figures 2-4.

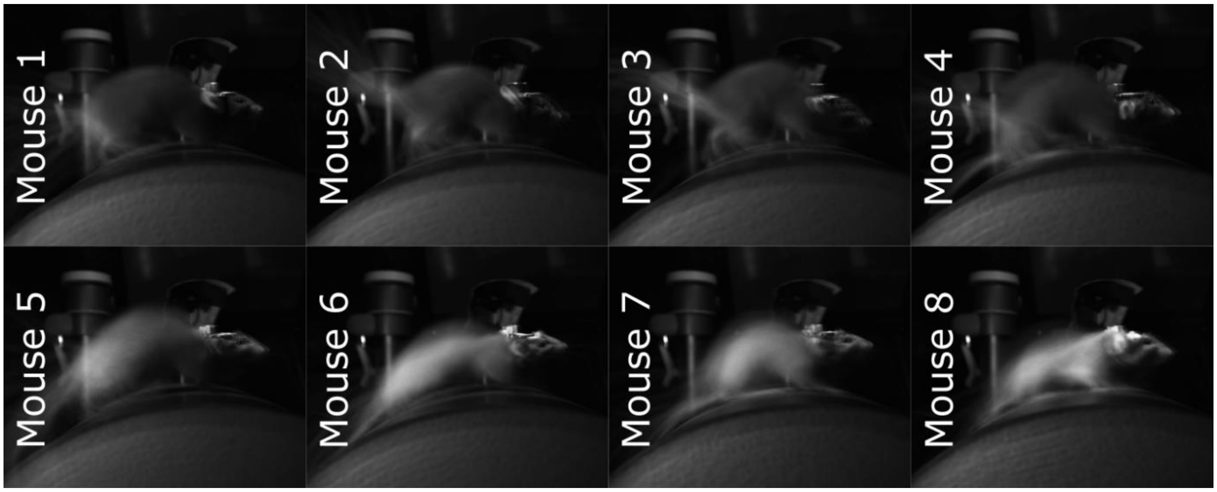


**Supplementary Figure 9.** *Superimposed first frames of each trial.* Randomly sampled superimposed frames for each recording session, for each mouse. The body posture changes, while the head position does not.

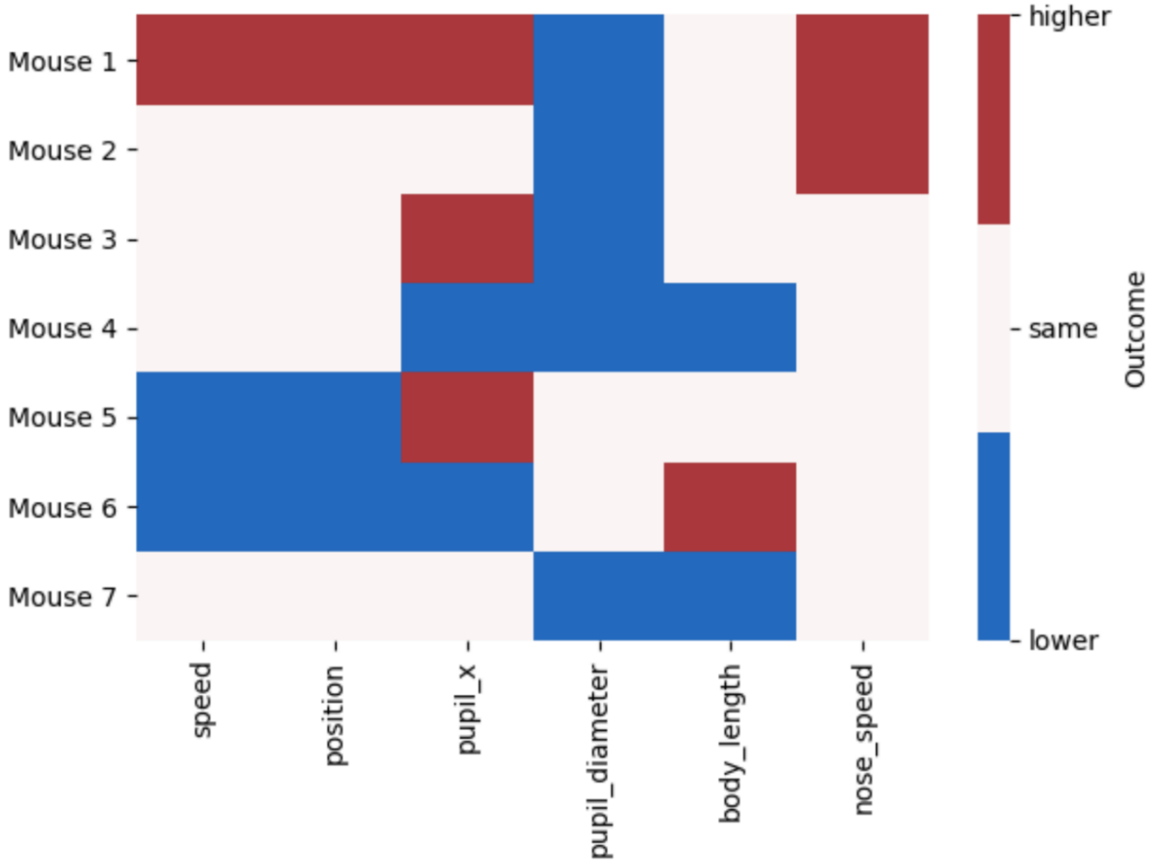


**Supplementary Figure 10.** *Summary of simulated foraging experiment results.* An overview mapping each mouse used in the simulated foraging paradigm to its outcome.

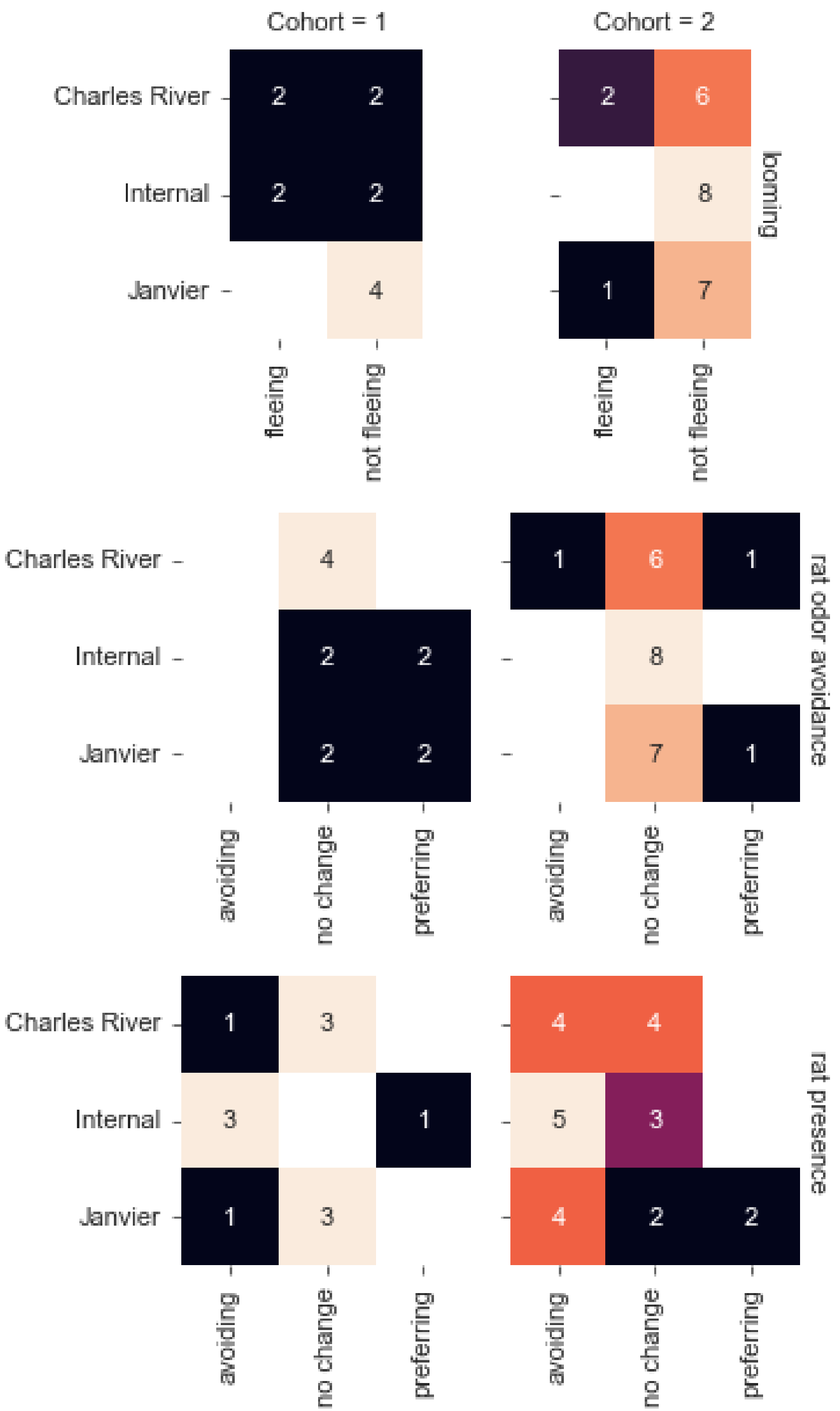


**Supplementary Figure 11.** *Summary of looming, rat odor & presence avoidance experiment results.* An overview mapping each mouse used in the verification experiments to its outcome. Grid rows are the 3 paradigms (looming, rat odor avoidance, and rat presence) and grid columns are the two cohorts. Each field consists of 3 rows, one per source, and 2-3 columns, based on the possible outcomes in the given paradigm. Color scale based on the number of mice.

### Extended Media

**Video 1.** Three simultaneous views of rat and mouse interaction from 3 cameras.

**Video 2.** Mouse position on treadmill.

**Video 3.** Pupil Position.

**Video 4.** Pupil size.

**Video 5.** Body posture.

**Video 6**. Nose movement.
